# Cultural variation impacts paternal and maternal genetic lineages of the Hmong-Mien and Sino-Tibetan groups from Thailand

**DOI:** 10.1101/2020.01.21.913582

**Authors:** Wibhu Kutanan, Rasmi Shoocongdej, Metawee Srikummool, Alexander Hübner, Thanatip Suttipai, Suparat Srithawong, Jatupol Kampuansai, Mark Stoneking

## Abstract

The Hmong-Mien (HM) and Sino-Tibetan (ST) speaking groups are known as hill tribes in Thailand; they were the subject of the first studies to show an impact of patrilocality vs. matrilocality on patterns of mitochondrial (mt) DNA vs. male-specific portion of the Y chromosome (MSY) variation. However, HM and ST groups have not been studied in as much detail; here we report and analyze 234 partial MSY sequences (∼2.3 mB) and 416 complete mtDNA sequences from 14 populations that, when combined with our previous published data, provides the largest dataset yet for the hill tribes. We find a striking difference between Hmong and IuMien (Mien-speaking) groups: the Hmong are genetically different from both the IuMien and all other Thai groups, whereas the IuMien are genetically more similar to other linguistic groups than to the Hmong. In general, we find less of an impact of patrilocality vs. matrilocality on patterns of mtDNA vs. MSY variation than previous studies. However, there is a dramatic difference in the frequency of MSY and mtDNA lineages of Northeast Asian (NEA) origin vs. Southeast Asian (SEA) origin in HM vs. ST groups: HM groups have high frequencies of NEA MSY lineages but lower frequencies of NEA mtDNA lineages, while ST groups show the opposite. A potential explanation is that the ancestors of Thai HM groups were patrilocal, while the ancestors of Thai ST groups were matrilocal. Overall, these results attest to the impact of cultural practices on patterns of mtDNA vs. MSY variation.

## Introduction

Thailand occupies the center of Mainland Southeast Asia (MSEA) and shares borders with several countries: Laos and Myanmar in the North and West, Laos in the Northeast, Cambodia in the East and Malaysia in the South (Fig. 1). With a census size of ∼68 million in 2015, there are 68 different recognized languages belonging to five different linguistic families: Tai-Kadai (TK, 90.5%), Austroasiatic (AA, 4.0%), Sino-Tibetan (ST, 3.2%), Austronesian (AN, 2.0%), and Hmong-Mien (HM, 0.3%) (Simons and Fennig 2018). Archaeological evidence indicates the presence of modern humans in the area of Thailand since the late Pleistocene (Shoocongdej, 2006; Higham, 2014). More recently, archaeogenetics studies indicate that modern AA-speaking groups in Southeast Asia (SEA) are descended from a dispersal of Neolithic farmers from southern China that occurred ∼4 thousand years ago (kya) (McColl et al., 2018; Lipson et al., 2018). Linguistic evidence supports the presence of AA people by at least 2.5 kya, while a later expansion of the TK speaking groups from southern China reached present-day Thailand around 2 kya (Blench, 2015; Bellwood 2018). And, historical evidence indicates that the homeland of many ST and HM speaking hill tribe groups (e.g., Akha, Lisu, Lahu, Karen, Hmong and IuMien) is in the area further north of Thailand, i.e. northern Myanmar, northern Laos and southern China, and most of these groups migrated to present-day Thailand ∼ 200 ya (Penth 2000; Schliesinger 2000). Thus, the present-day cultural and linguistic variation in Thailand has multiple sources, but the impact on genetic variation is still poorly understood.

**Figure 1.**
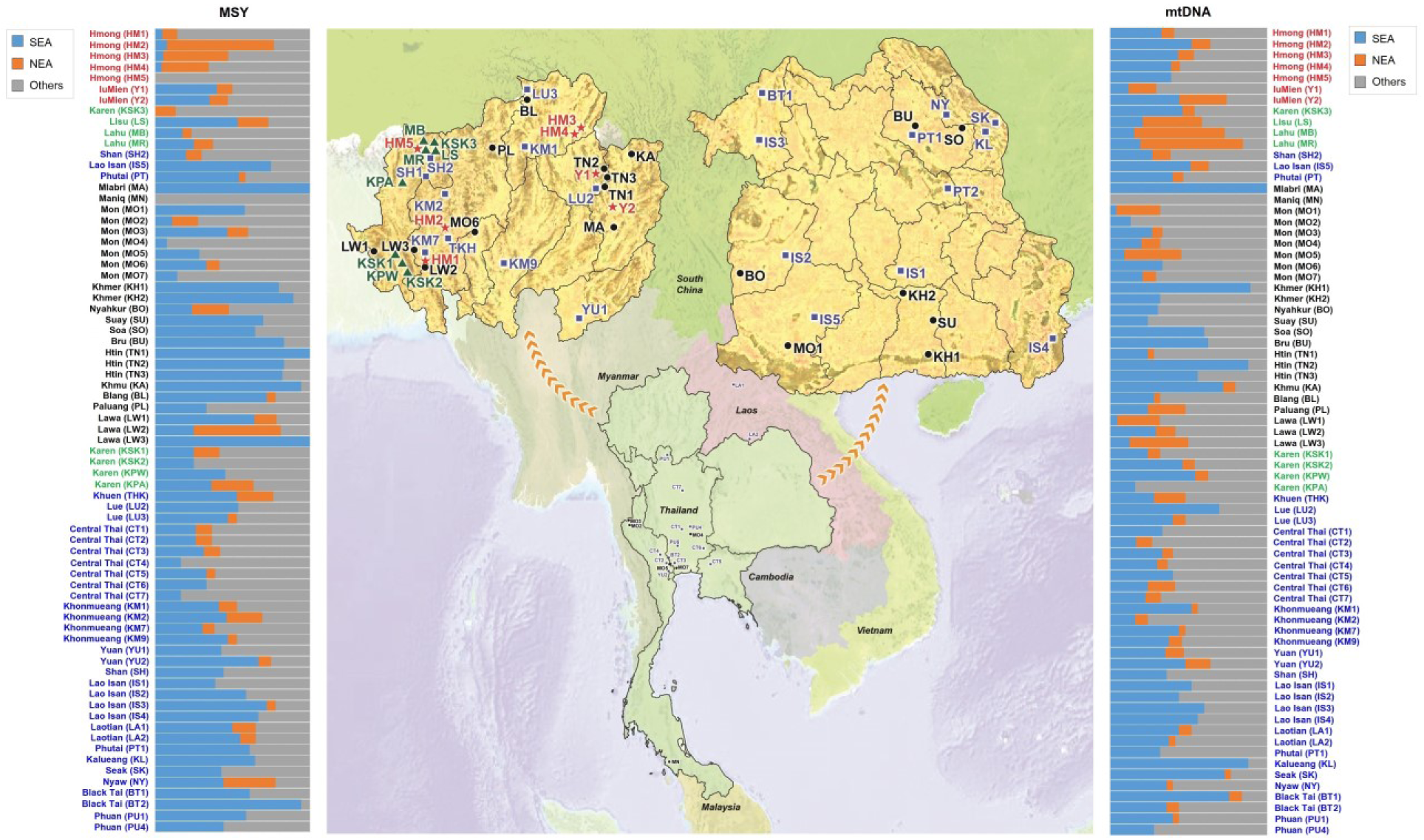
Map showing the location of the 14 populations sampled in the present study from northern Thailand (HM1-HM5, Y1-Y2, KSK3, MR, MR, LS and SH2) and northeastern Thailand (PT2 and IS5), together with 59 Thai/Lao populations sampled in previous studies (Kutanan et al., 2017; Kutanan et al., 2018a; Kutanan et al., 2018b; Kutanan et al., 2019). Red stars, green triangles, black circles and blue squares represent Hmong-Mien, Sino-Tibetan, Austroasiatic and Tai-Kadai speaking populations, respectively. The barplots on the left and right sides of the map depict the proportion of MSY and mtDNA haplogroups specific to Northeast Asia (NEA), Southeast Asia (SEA), or of unknown/other origin.

Based on geography, the Thai people can be characterized as highlanders vs. lowlanders. The hill tribes are highlanders, who inhabit the high mountainous northern and western region of Thailand. Consisting of ∼700,000 people, there are nine officially recognized hill tribes: the AA-speaking Lawa, Htin and Khmu; the HM-speaking Hmong and IuMien; and the ST-speaking Karen, Lahu (or Mussur), Akha and Lisu (Schliesinger 2000, 2001; Penth and Forbes 2004). In addition to living in a remote and isolated region of Thailand, the hill tribes are of interest for their cultural variation. In particular, post-marital residence pattern varies among the hill tribes, with some practicing patrilocality (i.e., following marriage, the woman moves to the residence of the man) while others are matrilocal (i.e., the man moves to the residence of the woman). The first study to document an effect of patrilocality vs. matrilocality on patterns of human mitochondrial (mt) DNA vs. male-specific portion of the Y chromosome (MSY) variation was carried out on the hill tribes (Oota et al. 2001), and has been further investigated in subsequent studies (Besaggio et al. 2007; Kutanan et al., 2019).

Our previous studies on the paternal and maternal genetic lineages and structure of many TK and AA groups indicated different and complex demographic histories in populations from Thailand and Laos (Kutanan et al., 2017; 2018a; 2018b; Kutanan et al., 2019). However, the ST and HM speaking groups have not been studied in as much detail. Here we generated and analyzed 416 complete mtDNA genome sequences and 234 partial sequences of the MSY from 14 populations belonging to 11 hill tribe HM and ST populations, and from three non-hill tribe TK populations: the Shan, who migrated recently from Myanmar and live in the mountainous area of northern Thailand; and the Phutai and Lao Isan from the Northeast of Thailand (Fig. 1). This is the first detailed genetic study of Thai HM speaking groups; unexpectedly, we find that the Hmong and IuMien groups have different histories, with the Hmong largely isolated from other groups and the IuMien having more in common with other groups. We also revisit the impact of patrilocality vs. matrilocality on patterns of mtDNA and MSY variation with higher-resolution methods and more populations than studied previously, and find less of a contrast in patterns of genetic variation between patrilocal and matrilocal groups than documented previously. However, we find a striking difference in Northeast Asian (NEA) ancestry vs. Southeast Asian (SEA) ancestry in HM vs. ST groups that we suggest may reflect differences in ancestral postmarital residence patterns.

## Results

### Genetic lineages

#### MSY

We combined the 234 newly generated sequences with 928 sequences from our previous studies (Kutanan et al., 2018a; Kutanan et al., 2019) for a total of 1,161sequences, of which 818 are distinct, from 73 populations; population details are listed in Table S1. The mean coverages of newly generated 234 MSY sequences of ∼2.3 mB range from 7x to 60x (overall average coverage 18x) (Table S2).

When combined with our previous Thai/Lao data, there are a total of 90 haplogroups identified (Table S3); 10 of these were not found in our previous studies. O1b1a1a (O-M95) (26.61%), O2a2b1a1 (O-Page23) (9.75%) and O1b1a1a1a1 (O-M88) (7.15%) are prevalent in almost all groups (overall frequency 43.52%) (Fig. S1). Haplogroup O2a2a1a2a1a2 (O-N5) and C-F845 are mostly prevalent in HM groups while haplogroup F is the dominant haplogroup of the Lahu (MR and MB). The coalescent ages of these three haplogroups are ∼2.45 kya (HPD: 2.88-1.13 kya) for O2a2a1a2a1a2 (O-N5), ∼12.54 kya (HPD: 17.16-4.09 kya) for C- F845 and ∼16.00 kya (HPD: 22.59-12.26 kya) for haplogroup F. However, if we focus on HM or Lahu clades of the MCC tree of haplogroup C-F845, the age is ∼2.85 kya (HPD: 4.25-1.08 kya) and ∼0.58 ya (HPD: 1.55-0.29) for haplogroup F (Fig. S2).

When we focus on the MSY lineages that are prevalent in Northeast Asia (NEA), i.e. C2e*, D-M174 and N* in our samples, the frequency of NEA lineages is greater than 30% among the Hmong (HM2-HM4), Lawa (LW2), and Karen (KPA) from northern Thailand, and the Nyaw (NY) from northeastern Thailand. In contrast O1b*, which is the predominant lineage in Southeast Asia (SEA) and at high frequency in AA speaking people (Cai et al., 2011; Kutanan et al., 2019), is the major lineage in the Thai/Lao AA speaking group (except for some Mon populations who show evidence of admixture with Central Thai populations). Interestingly, there is heterogeneity in the frequency of NEA/SEA lineages in the Hmong and Lawa groups: among the five Hmong populations, HM5 completely lacks NEA lineages, while within the Lawa groups, LW2 has 56% NEA lineages while LW3 has exclusively SEA lineages (Fig. 1).

#### mtDNA

We generated 416 complete mtDNA sequences with mean coverage ranging from 35x to 7752x (overall average coverage 1934x) (Table S4). When combined with 1,434 sequences from our previous studies (Kutanan et al., 2017; Kutanan et al., 2018a; Kutanan et al., 2018b) there are in total 1,850 sequences belonging to 73 populations, with 1,125 haplotypes. When combined with our previous data there are a total of 285 haplogroups (Table S5); several were not reported previously from Thai/Lao populations and these are specific to some populations, e.g. B4a5 (specific to the Hmong), and B5a1c1a, B5a1c1a1, D4e1a3 and F1g1 (specific to HM groups).

The coalescent ages (Fig. S3) of the prevalent lineages of HM groups are ∼10.67 kya (HPD: 11.27-3.31kya) for B5a1c1a*, ∼1.53 kya (HPD: 3.96- 0.94 kya) for B5a1c1a1, ∼6.83 kya (HPD: 7.09-1.55 kya) for D4e1a3, ∼11.54 kya (HPD: 16.04-5.84 kya) for F1g1. B4a5, specific for the Hmong, has a coalescent age of ∼6.63 (HPD: 11.16-3.22 kya). The coalescent ages of D4j1a1 and G1c, abundant in the Lahu, are ∼9.23 kya (HPD: 12.07-4.85 kya) and ∼3.88 kya (HPD: 4.70-0.21 kya). In addition, the Lahu-specific clade of D4j1a1 sequences is dated to 2.49 kya (HPD: 5.83-1.53 kya). The coalescent age of B6a1a, abundant in the Karen, is ∼6.69 kya (HPD: 11.62-4.25 kya).

Haplogroup A*, D* and G* are predominant in NEA populations and we find that the frequency of these NEA lineages is greater than 30% in Lahu (MB and MR), Lawa (LW3), Lisu (LS), and IuMien (Y2) from northern Thailand, and in Mon (MO5) from central Thailand. In contrast, the predominant SEA haplogroups (B5*, F1a*, M7b* and R9b*) are at highest frequency in TK and AA speaking groups, indicating genetic similarity between these two groups. Strikingly, the populations with the highest frequencies of NEA MSY lineages are not the same as the populations with the highest frequencies of NEA mtDNA lineages (Fig. 1), which we suggest below may reflect differences in ancestral postmarital residence patterns.

### Genetic diversity

#### MSY

In general, the HM, AA and ST groups tend to have lower genetic diversity values than the TK groups (Table S6). By contrast, genetic diversities of the HM groups are not statistically different from the AA and ST groups, nor do the ST and AA groups differ significantly in genetic diversity values (Table S6). At the individual population level, out of 63 populations, the Hmong (HM1) shows lower haplotype diversity than all other groups except for two hunter-gatherer groups (Mlabri (MA) and Maniq (MN)) and the Htin Mal (TN1). HM1 and HM5 have lower genetic diversity than the other HM populations, although HM2 shows the highest MPD values. Generally, the Hmong (HM1-HM5) groups show lower haplotype and haplogroup diversity than the In Mien (Y1-Y2) (Fig. 2A and 2B). Of the eight ST speaking populations, the newly-studied Karen group (KSK3) exhibits lower genetic diversities while the Lisu (LS) and KSK1 have higher genetic diversities than the other ST speaking populations (Fig. 2A-2D). Although low genetic diversities are observed in HM1, a significantly low Tajima’s *D* value (Fig. 2D) suggests recent paternal expansion in this group. Significant negative Tajima’s *D* values were observed more frequently in the TK than in the AA and HM groups (*P* < 0.05: 11/34 for TK, 6/24 for AA and 2/7 for HM) but no significant Tajima’s *D* values were observed in any of the ST-speaking groups.

**Figure 2.**
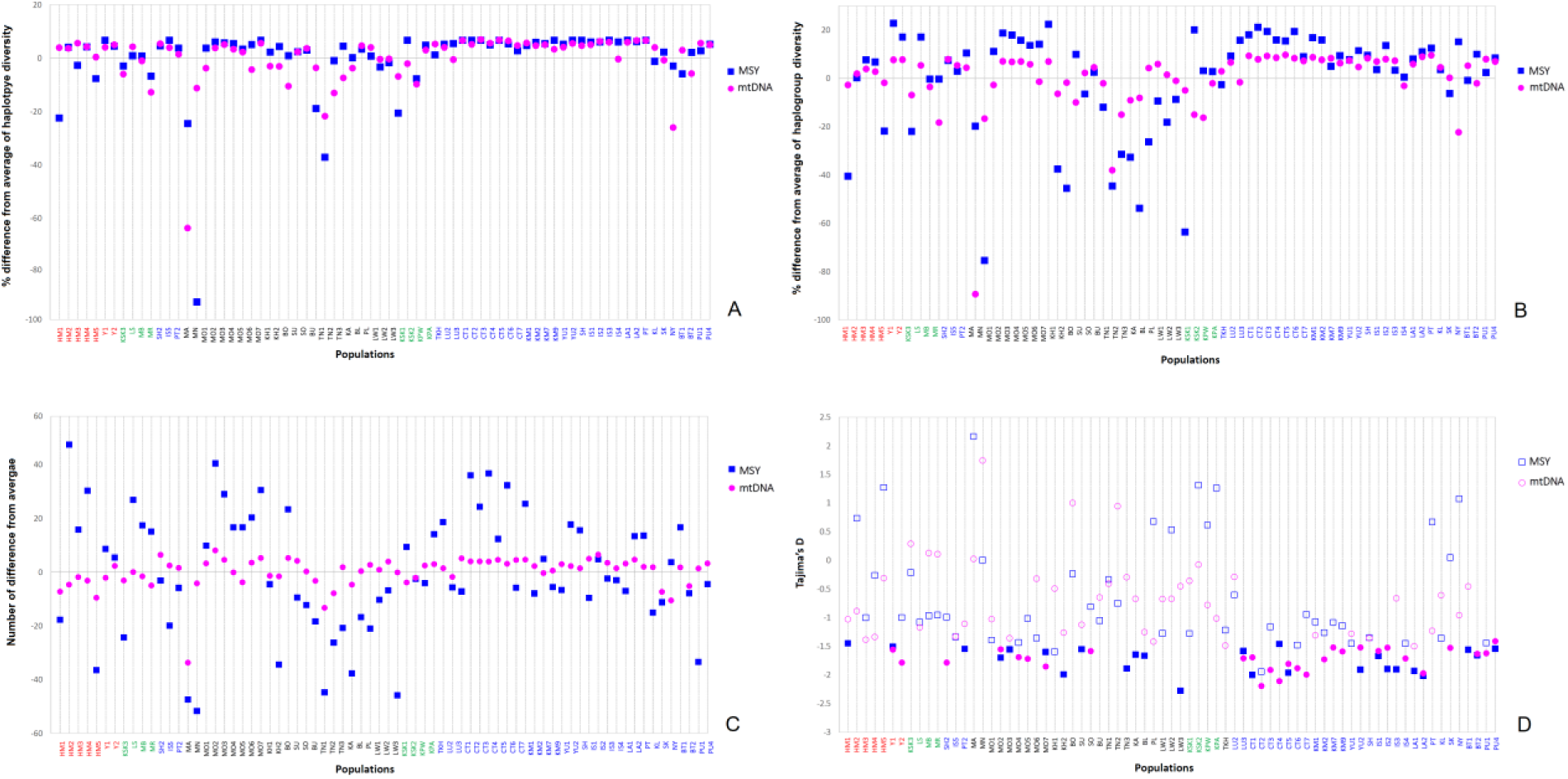
Genetic diversity values for the MSY and mtDNA in 73 populations, shown as the percent difference from the average of: (*A*) haplotype diversity, (*B*) haplogroup diversity and (*C*) MPD. The grey line shows the mean across populations. (*D*) Tajima’s *D* values; solid symbols indicate values significantly different from zero (*P* < 0.05). More information and all genetic diversity values are provided in Table S1. The new populations are placed at the left of the figure. Population names are color-coded according to language family; red, green, black and blue represent HM, ST, AA and TK speaking populations, respectively.

#### mtDNA

Along with the Mlabri (MA), Htin (TN1 and TN2) and Seak (SK), the ST speaking Lahu or Mussur (MR) shows low mtDNA haplotype and haplogroup diversities whereas the Lisu (LS) shows higher genetic diversities than the other ST populations (Fig. 2A-2C). In contrast to the MSY, the Hmong groups exhibited generally higher genetic diversities than the ST and AA speaking groups (Table S6). Both the ST and AA groups have lower genetic diversity values than the TK groups (Table S6). As also seen in the MSY results, both IuMien populations (Y1 and Y2) show higher genetic diversities than the other Hmong and ST speaking groups (Fig. 2A-2C). In agreement with the MSY results, a significantly negative Tajima’s *D* value was observed more frequently for the TK than for the AA and HM groups (*P* < 0.05: 21/34 for TK, 5/24 for AA and 2/7 for HM). Interestingly, the ST-speaking groups show no significant Tajima’s *D* values while the two IuMien groups both show significant negative Tajima’s *D* values (Fig. 2D).

### The Analysis of Molecular Variance (AMOVA)

#### MSY

The AMOVA results indicate that the variation among populations accounts for 13.72% of the total MSY genetic variance (Table 1). The HM group shows the greatest genetic heterogeneity among populations, followed by the AA and ST groups; the TK group shows the lowest among-population variation. The Thai Hmong, with five populations sampled, shows higher variation among the populations than do the other hill tribe groups. When HM and ST populations from Vietnam (Macholt et al., 2109) were included in the analysis, genetic variation among populations of the ST and HM groups increased substantially, suggesting some differentiation between Vietnamese and Thai populations belonging to these two groups. However, a direct comparison of Thai ST vs. Vietnamese ST, and Thai HM vs. Vietnamese HM groups, showed no significant differences between groups. The variation among populations within groups of the IuMien and Lahu were much lower than for the Hmong, indicating genetic heterogeneity of the Hmong and ST populations and more homogeneity for the IuMien and Lahu. The MSY genetic variation showed significant differences among the four language families (HM, ST, AA and TK), but the variation among groups was lower than the variation among populations within each group, indicating that language families do not correspond to genetic structure. All pairwise comparisons of the four language families showed significant differences among groups, but these were on the same order as the differences among populations within the same group. However, when the HM were separated into Hmong and IuMien, the pairwise comparisons of Hmong with other language families remained significant, while for the IuMien there were no significant differences with other language families, suggesting some differences between Hmong and Mien groups. The lowest variation was between the TK and AA groups, indicating a relatively close genetic relationship between these two.

**Table 1.** AMOVA results.

| Groups | a | b | Percent variation |  |  |  |  |  |
| --- | --- | --- | --- | --- | --- | --- | --- | --- |
|  |  |  | Within populations |  | Between populations within groups |  | Among groups |  |
|  |  |  | MSY | mtDNA | MSY | mtDNA | MSY | mtDNA |
| Total | 1 | 73<br>(71) | 86.28<br>(86.85) | 91.54<br>(92.33) | 13.72**<br>(13.15**) | 8.46**<br>(7.67**) |  |  |
| Hmong-Mien (HM) | 1 | 7 | 81.83 | 96.6 | 18.17** | 3.4** |  |  |
| Austroasiatic (AA) | 1 | 24<br>(22) | 82.15<br>(83.92) | 85.96<br>(88.47) | 17.85**<br>(16.08**) | 14.04**<br>(11.53**) |  |  |
| Sino-Tibetan (ST) | 1 | 8 | 87 | 89.76 | 13** | 10.24** |  |  |
| Tai-Kadai (TK) | 1 | 34 | 95.59** | 95.86 | 4.41** | 4.14** |  |  |
| HM<br>(Thailand+Vietnam) | 1 | 11 | 82 | 94.35 | 18** | 5.65** |  |  |
| ST (Thailand+Vietnam) | 1 | 12 | 74.96 | 87.64 | 25.04** | 12.36** |  |  |
| Patrilocal | 1 | 13 | 72.57** | 93.64** | 27.43** | 6.36** |  |  |
| Matrilocal | 1 | 10 | 75.51** | 83.54** | 24.49** | 16.46** |  |  |
| Hmong | 1 | 5 | 81.87 | 99.47 | 18.13** | 0.53 |  |  |
| Karen | 1 | 5 | 90.89 | 94.15 | 9.11** | 5.85** |  |  |
| Mon | 1 | 7 | 96.5 | 93.1 | 3.5** | 6.90* |  |  |
| H'tin | 1 | 3 | 90.11 | 74.29 | 9.89** | 25.71* |  |  |
| Lawa | 1 | 3 | 64.35 | 92.22 | 35.65** | 7.78* |  |  |
| Hmong<br>(Thailand+Vietnam) | 1 | 6 | 78.75 | 98.92 | 21.25** | 1.08 |  |  |
| IuMien<br>(Thailand+Vietnam) | 1 | 3 | 94.42 | 95.5 | 5.58** | 4.5** |  |  |
| Lahu<br>(Thailand+Vietnam) | 1 | 3 | 95.12 | 86.53 | 4.88* | 13.47** |  |  |
| Language (AA, TK,<br>HM, ST) | 4 | 73<br>(71) | 84.69**<br>(85.26**) | 91.21**<br>(91.95**) | 10.10**<br>(9.58**) | 7.7**<br>(6.85**) | 5.21**<br>(5.16**) | 1.09**<br>(1.20**) |
| HM vs. AA | 2 | 31<br>(29) | 70.24**<br>(71.52**) | 86.92**<br>(88.46**) | 15.18**<br>(14.17**) | 11.56**<br>(9.53**) | 14.58**<br>(14.31**) | 1.52*<br>(2.02**) |
| HM vs. ST | 2 | 15 | 78.13** | 89.23** | 14.59** | 6.69** | 7.28** | 4.09** |
| HM vs. TK | 2 | 41 | 82.85** | 94.3** | 6.11** | 4.00** | 11.04** | 1.70** |
| Hmong vs. AA | 2 | 29<br>(27) | 65.55**<br>(66.86**) | 85.44**<br>(87.02**) | 14.06**<br>(13.04**) | 11.79**<br>(9.59**) | 20.39**<br>(20.10**) | 2.77**<br>(3.40**) |
| Hmong vs. ST | 2 | 13 | 75.18** | 87.78** | 13.52** | 6.38** | 11.30** | 5.84** |
| Hmong vs. TK | 2 | 39 | 78.04** | 93.20** | 5.19** | 3.72** | 16.77** | 3.08** |
| IuMien vs. AA | 2 | 26<br>(24) | 83.01**<br>(84.54**) | 89.00**<br>(90.68**) | 16.92**<br>(15.21**) | 13.97**<br>(11.41**) | 0.07<br>(0.26) | -2.97<br>(-2.09) |
| IuMien vs. ST | 2 | 10 | 88.45** | 90.69** | 11.46** | 9.35** | 0.09 | -0.04 |
| IuMien vs. TK | 2 | 36 | 95.92** | 95.93** | 4.28** | 4.14** | -0.19 | -0.08 |
| Thai HM vs. Vietnam<br>HM | 2 | 11 | 80.82** | 93.86** | 15.98** | 5.01** | 3.2 | 1.13 |
| AA vs. ST | 2 | 30 | 78.04**<br>(79.40**) | 86.02**<br>(87.98**) | 15.54**<br>(14.33**) | 13.03**<br>(11.13**) | 6.41**<br>(6.27**) | 0.95*<br>(0.9) |
| AA vs. TK | 2 | 58<br>(56) | 89.71**<br>(90.51**) | 91.87**<br>(92.83**) | 9.27**<br>(8.57**) | 7.92**<br>(6.87**) | 1.02*<br>(0.92*) | 0.21<br>(0.3*) |
| ST vs. TK | 2 | 42 | 90.22** | 92.94** | 5.56** | 5.02** | 4.22** | 2.05** |
| Thai ST vs. Vietnam ST | 2 | 12 | 74.65** | 86.85** | 24.44** | 11.23** | 0.91 | 1.92 |
| ** indicate $P < 0.01$, * indicates $P < 0.05$<br>a = number of groups; b = number of populations; numbers/values in parentheses were calculated by excluding the two hunter-gather groups, Mlabri (MA) and Maniq (MN) | | | | | | | | |

#### mtDNA

The total mtDNA variation among populations of 8.46% was lower than for the MSY (Table 1). The mtDNA variation for the AA, ST and TK groups was about the same as for the MSY, but was substantially less for the HM. The Htin have by far the largest variation among populations, while the Hmong show non-significant mtDNA variation among populations (0.53%), reflecting genetic homogeneity in their maternal side. The mtDNA genetic variation among the four language families (HM, ST, AA and TK) was much smaller (1.09%) than the variation among populations assigned to each group (7.7%), indicating that as with the MSY, language families do not correspond to the genetic structure of these populations. The variation between pairs of linguistic groups shows in all comparisons that the variation between groups is lower than the variation among populations within groups. As with the MSY, the lowest variation (which is not significantly different from zero) is between the TK and AA groups, further supporting a close relationship between these two groups in Thailand.

The pooled mtDNA data of Lahu from Vietnam and Thailand revealed much larger variation for mtDNA (13.47%) than for the MSY (4.88%), in contrast to the larger MSY than mtDNA variation observed when pooling data from other groups from Thailand and Vietnam. In particular, the mtDNA variation among Hmong groups from Thailand and Vietnam was only 1.08%, which is not significantly different from zero. When the Hmong and IuMien were separately compared with other linguistic groups, significant differences were observed for the Hmong but not for IuMien, similar to the MSY results and further supporting the difference between Hmong and Mien groups.

### Population Affinity

#### MSY

Shared haplotypes within populations are an indication of smaller population size and increase relatedness among individuals, while shared haplotypes between populations are an indication of recent shared ancestry or contact. There were shared MSY haplotypes within the HM groups, and some sharing between them and a few TK-speaking groups, except for HM5 and Y1, who did not share any haplotypes with any other populations (Fig. 3A). The Lisu shared haplotypes with both Lahu (MB and MR) populations, while both Lahu populations shared haplotypes among themselves and also with one group of central Thai (CT5). The newly studied Karen (KSK3) shared haplotypes with the other Karen populations (KSK1, KSK2 and KPW), and also with their neighbors, i.e. Shan (SH2) and Lawa (LW1) (Fig. 3A).

**Figure 3.**
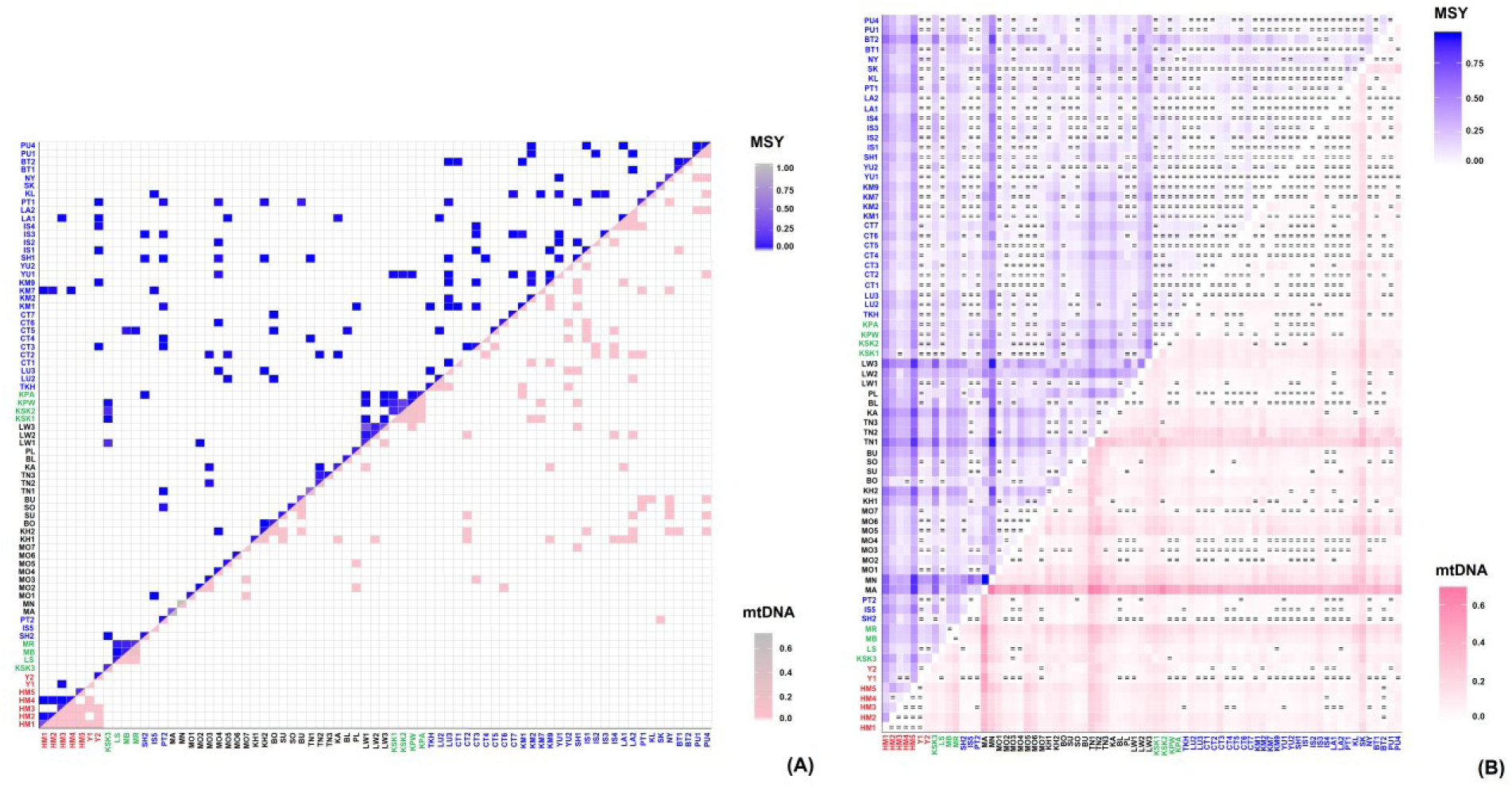
(*A*) Frequency of shared MSY (above diagonal) and mtDNA (below diagonal) haplotypes within and between populations. (*B*) heat plot of *Φ*_st_ values based on MSY (above diagonal) and mtDNA (below diagonal) haplotypes. The “=” symbol indicates *Φ*_st_ values that are not significantly different from zero (*P* > 0.05). The new populations are placed at the left of the figure. Population names are color-coded according to language family; red, green, black and blue represent HM, ST, AA and TK speaking populations, respectively.

Genetic distance values are a further indication of genetic relationships among populations; the genetic distances (*Φ*_st_ values) indicate, in general, genetic heterogeneity among AA populations and homogeneity among TK populations, as well as genetic differences between the AA (except the Mon) and TK populations. For the newly-studied HM groups, significant genetic differences between the Hmong and almost all other populations were observed, whereas the *Φ*_st_ values for comparisons of the IuMien (Y1 and Y2) and Lisu with many populations were not significant (Fig. 3B). The new Karen group is significantly different from almost all populations except SH2 and KSK1. The two Lahu populations are genetically distinct from all other populations (except each other).

To further visualize the relationships based on the *Φ*_st_ distance matrix, we carried out an MDS analysis. The MDS plot for three dimensions indicates genetic distinction of the Maniq (MN), the hunter-gatherer group from southern Thailand, the Hmong groups (HM1-HM5) and the Karen (KSK3) (Fig. S4), as further indicated in the MDS heat plot (Fig. S5). Based on the MDS results for both the MSY and mtDNA, we removed five highly-diverged populations (MA, MN, TN1, TN2 and SK); a three-dimension MDS for the remaining 68 populations has an acceptable stress value (Fig. 4A-4C). There was overall some clustering of populations according to language family, albeit with some overlapping between them. The Hmong populations are quite distinct from all other groups, whereas the IuMien populations are more similar to other groups than to the Hmong groups. The TK overlap with AA groups, but the AA are more spread out, indicating more genetic divergence of AA groups. KSK3 and, to a lesser extent, KSK2 and both Lahu populations are distinct from the other ST groups.

**Figure 4.**
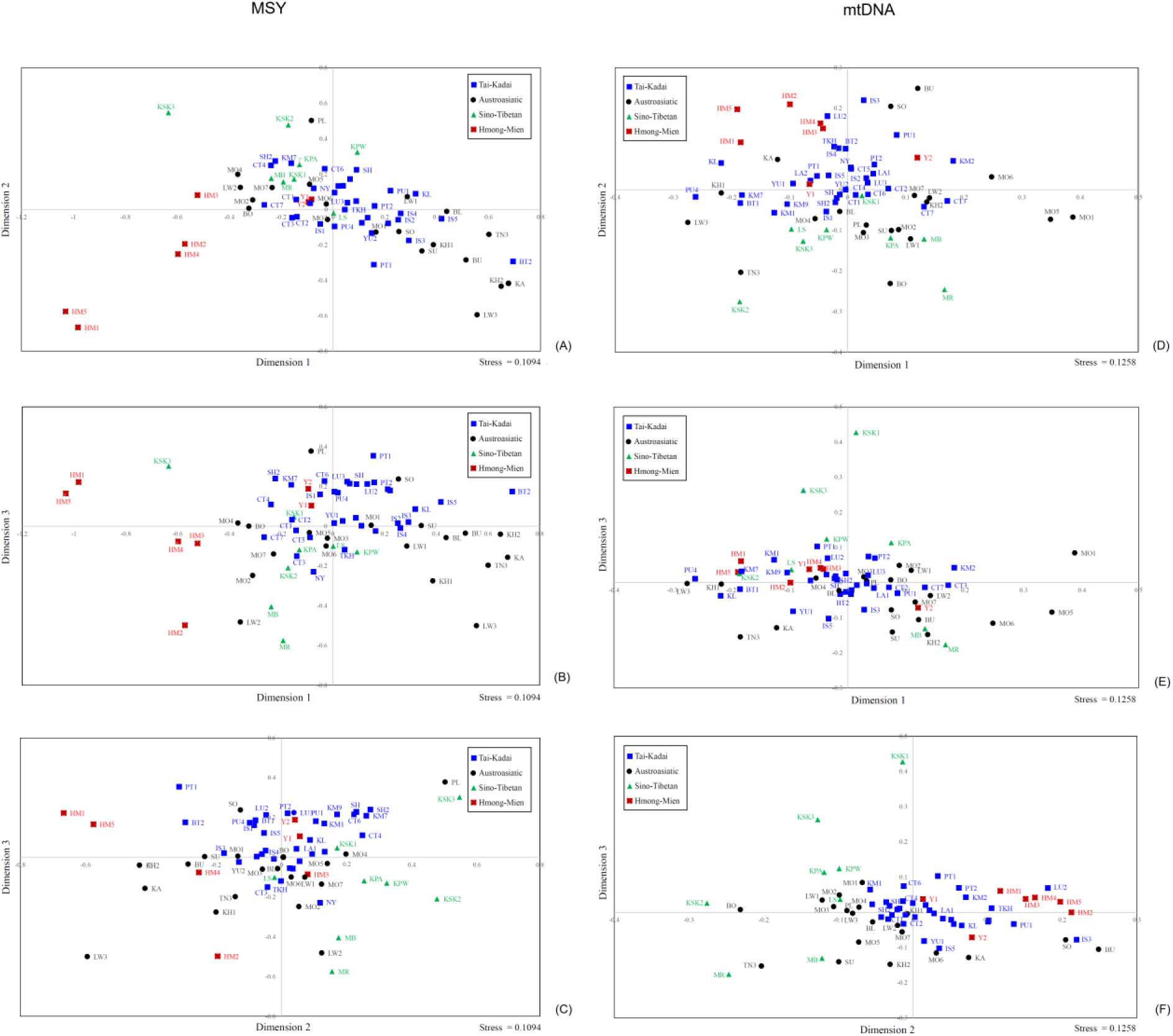
The three-dimensional MDS plot based on the *Φ*_st_ distance matrix for 68 Thai/Lao populations (after removal of Maniq, Mlabri, Htin (TN1, TN2) and Seak (SK)) for (A) MSY and (B) mtDNA. The stress values are 0.1094 for MSY and 0.1258 for mtDNA.

To investigate the relationships of Thai/Lao populations with other SEA populations, we included available comparable sequencing data from populations from Vietnam, southern China, and Myanmar. The MDS plot based on the *Φst* distance matrix of 88 populations (Fig. 5A-5C; the same outliers are excluded as in Fig. 4A-4C) shows clustering of the Vietnamese HM-speaking Pathen and Hmong with the Thail Hmong populations, while the Vietnamese HM- speaking Yao and Hani are more similar to the Thai IuMien groups. The Karen (KSK3) remains distinct, but shows a close genetic relatedness to Burmese. Some of the TK speaking groups from Vietnam, i.e. Nung, Tay and Thai, are close to the TK populations from northeastern Thailand, e.g. Phutai (PT1 and PT2), Kalueang (KL) and Black Tai (BT1).

**Figure 5.**
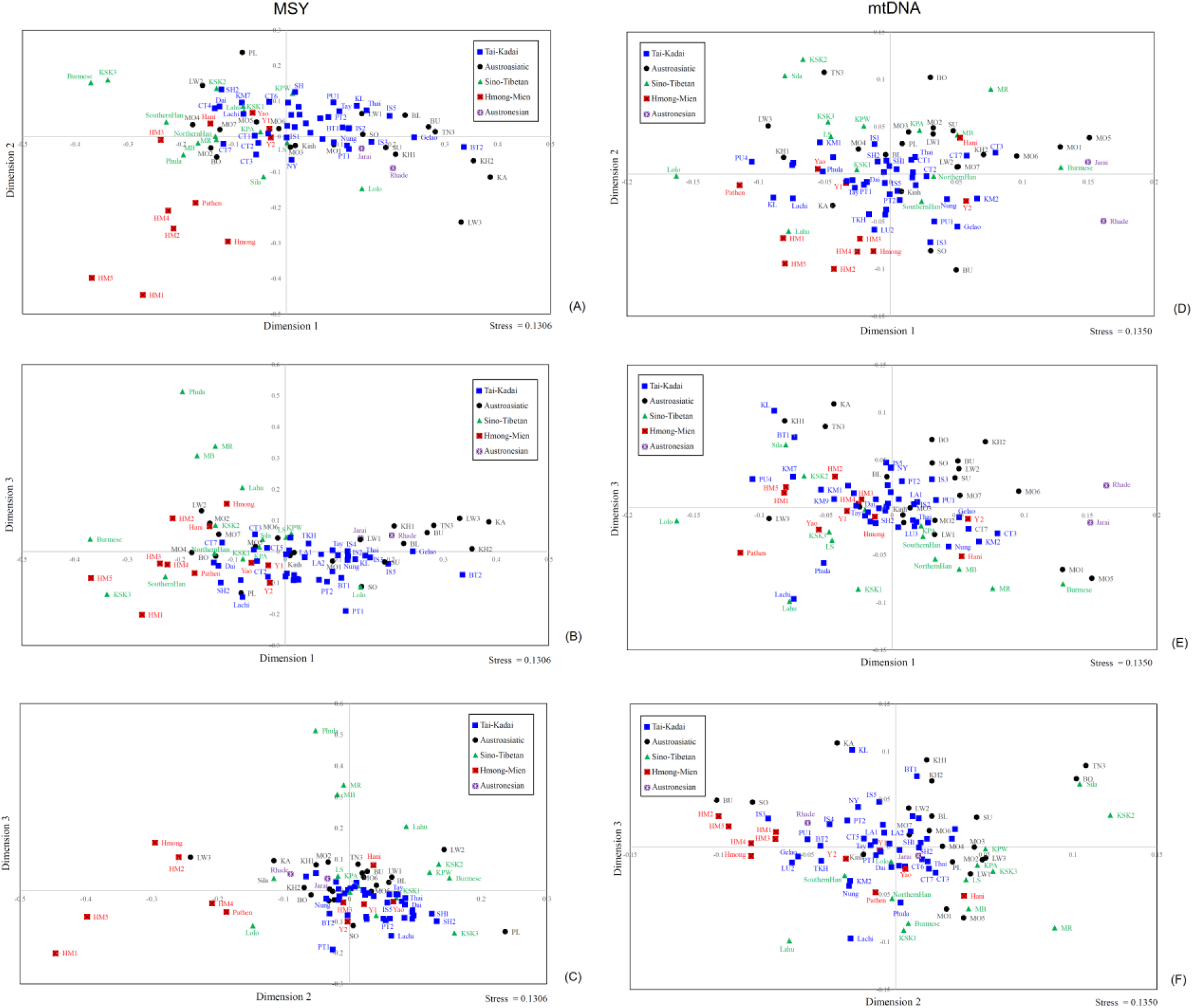
The two-dimensional MDS plot based on the *Φ*_st_ distance matrix for 88 SEA populations. (A) MSY and (B) mtDNA. The stress values are 0.1306 for MSY and 0.1350 for mtDNA.

To investigate which MSY haplogroups might be driving population relationships, we carried out a Correspondence Analysis (CA), which is based on haplogroup frequencies. The results (Fig. S6) indicate that the genetic distinctiveness of the Hmong reflects high frequencies of haplogroups O2a2a1a2a1a2 (O-N5) and C2e2 (C-F845). One IuMein population (Y1) is positioned between the Hmong and other Thai/Lao populations, reflecting haplogroups D (D- M174) and C2e1b1 (C-F1319). The second dimension distinguishes the two Lahu groups (MB, MR) and Seak (SK), based on haplogroup F (F-M89). The Soa (SO) occupy an intermediate position, based on O1b1a1a1a2a1a (O-Z24091). The third dimension distinguishes three of the Karen populations (KSK1, KSK2 and KPW), based on haplogroups O1b1a1a1b1a1 (O- FGC29907) and G1 (G-M342). Further dimensions distinguish an AA group (LW2) based on haplogroups O2a2b2a2 (O-F706) and N (N-M231), while the AA group MO2 and TK group CT7 are distinguished based on haplogroups R1a1a1b2a1b (R-Y6) and J2a1 (J-L26).

#### mtDNA

With respect to mtDNA haplotype sharing (Fig. 3A), the HM populations (HM1- HM5) share mtDNA haplotypes extensively with each other, including with the IuMien populations (Y1 and Y2), but do not share haplotypes with any other population, reflecting their unique genetic structure. As with the MSY results, the Lisu (LS) only shares haplotypes with both Lahu populations (MB and MR), indicating contact between them. The newly studied Karen population (KSK3) also exhibits large differences from the other Karen groups: there is no mtDNA haplotype sharing between the KSK3 and the other Karen populations, while some of the other Karen populations do share haplotypes with one another.

The heat plot of *Φst* genetic distances (Fig. 3B) also supports genetic distinction of the HM from other Thai/Lao populations, and mostly nonsignificant *Φ*_st_ values among them, with the exception of the IuMien populations and HM3, who show more similarity to other Thai/Lao populations. However, consistent with the MSY results, Lisu are not significantly different from several TK, AA and ST speaking populations, while the two Lahu populations do not differ significantly from each other, but do show significant differentiation from the other Thai/Lao groups.

The MDS plots based on *Φ*_st_ values for the Thai/Lao populations show greatest genetic divergence for the MA, the hunter gatherer group from northern Thailand, followed by their linguistic relatives, the Htin (TN1, TN2) and Seak (SK) (Fig. S4). After removal of the same five outliers as for the MSY analysis (MA, MN, TN1, TN2, and SK), the MDS analysis based on dimensions 1 and 2 shows separation between the Hmong populations and the ST populations, with the IuMien populations rather closer to the cloud of TK populations around the center of the plot. The Karen groups are further differentiated by dimension 3 (Fig. 4D-4F).

The MDS plot based on the *Φ*_st_ distance matrix that includes comparative data from other SEA populations (Fig. 5D-5F) shows that the HM speaking populations from Thailand and Vietnam tend to cluster together, except the Thai/Vietnam IuMen and Hani are closer to other populations, consistent with the MSY results. The Vietnamese Lahu are quite distinct from the Thai Lahu, and in fact are closer to the Thai HM groups. Interestingly, the ST speaking populations are about as heterogeneous as the AA speaking groups.

The CA analysis based on mtDNA haplogroup frequencies (Fig. S7) further confirms the distinctiveness of the HM groups based on several haplogroups, i.e. B5a1c1a, B5a1c1a1, B4a5, C7a, D4e1a3, F1g1, F1g2, N9a10 (16311C) and M74a . The Lahu and MO5 were distinguished in the third dimension, reflecting haplogroups B4e and D4j1a1. In the fourth dimension several groups are distinguished via many specific lineages, including all of the Karen groups and two Mon groups from the border between Thailand and Myanmar.

### Bayesian Skyline Plots

#### MSY

The Bayesian Skyline Plots (BSPs) of population size change (*N*e) over time were constructed for each ethnicity. For the MSY, different trends were observed for different groups (Fig. 6). The *N*e of the Hmong gradually increased since ∼30 kya and then declined ∼2-3 kya, while for the Lahu the *N*e remained stable for a long period of time and then was sharply reduced around ∼1 kya. The Karen, Shan and Phutai showed a similar trend: the *N*e gradually increased, and then decreased ∼5 kya, with sharp increases ∼2-3 kya, followed by another decrease ∼1 kya. The *N*e for the IuMien slightly increased, and then decreased ∼2-3 kya.

**Figure 6.**
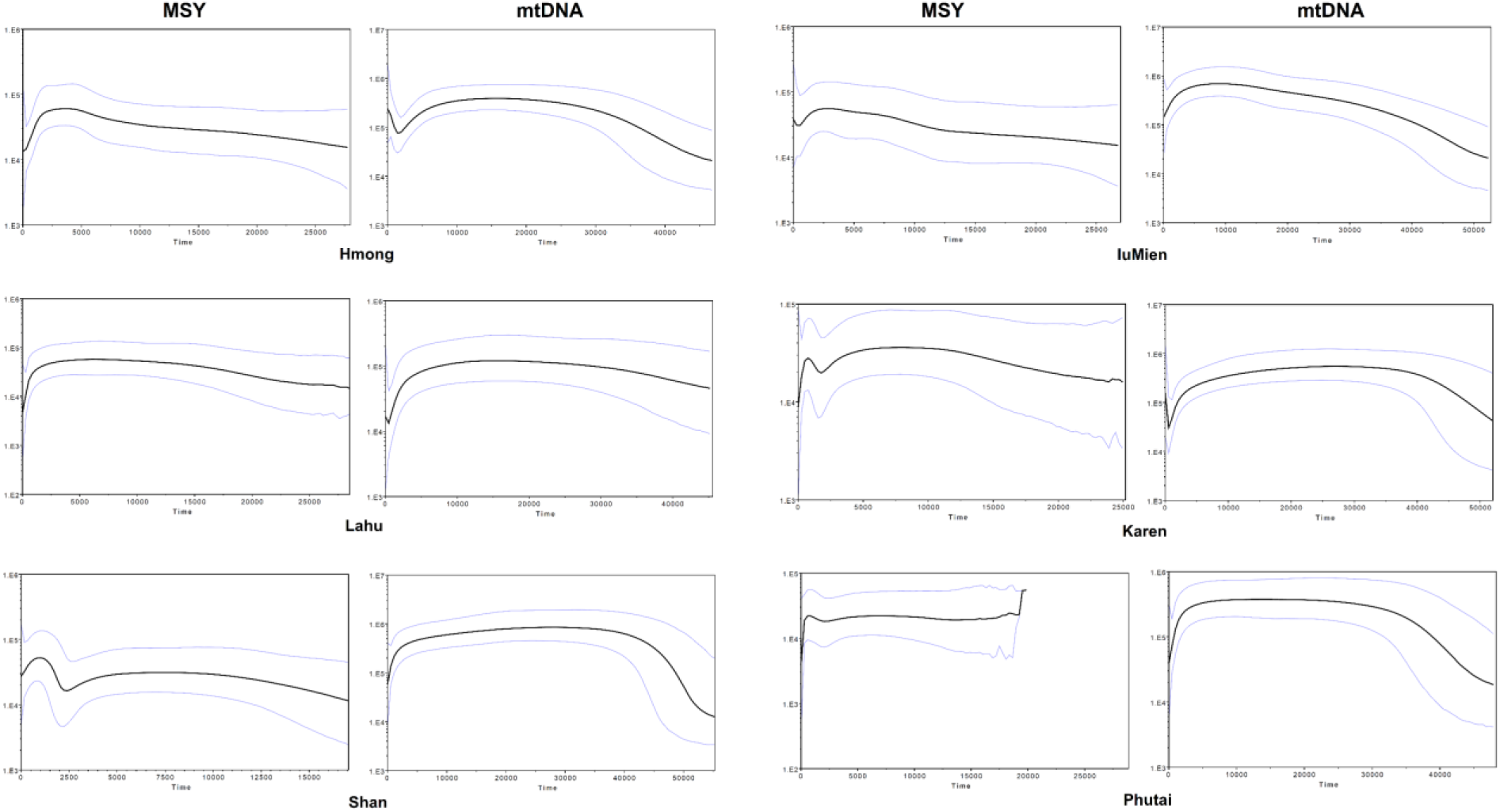
The BSPs based on the MSY and mtDNA for the Hmong, IuMien, Lahu, Karen, Shan and Phutai groups. Solid lines are the median estimated effective population size (y axis) through time from the present in years (x axis). The 95% highest posterior density limits are indicated by light-purple lines.

#### mtDNA

The BSPs for each ethnicity show that several groups, i.e. Lahu, Shan, Phutai, show a common trend of *N*e increasing 30–50 kya, and then stable until a decline ∼2–5 kya (Fig. 6; BSPs for each individual population are in Fig. S8). However, the Hmong and Karen showed a different pattern, namely a decrease in *N*e ∼5 kya followed by rapid growth ∼2.0–2.5 kya for the Hmong, and ∼1.0 kya for the Karen.

### Patrilocal vs. Matrilocal genetic variation

There are nine official hilltribes in Thailand: the AA-speaking Lawa, Htin, and Khmu; the HM-speaking Hmong and IuMien; and the ST-speaking Karen, Lahu, Lisu and Akha. The Lahu, Karen, and Htin are matrilocal (i.e., the husband moves to the residence of the wife after marriage) whereas the others are patrilocal. Our previous study (Kutanan et al., 2019) had investigated four hill tribes (Lawa, Htin, Khmu ad Karen); here we add data from four additional hill tribe groups (Hmong, IuMien, Lahu, and Lisu) for a total of 23 populations belonging to eight hill tribes. Moreover, although the Palaung is not officially recognized as a hill tribe group, we include them in the analysis because they are minority people come from the same mountainous region of northern Thailand.

The Hmong (HM1-HM5), IuMien (Y1-Y2), Lisu (LS), Khmu (KA), Lawa (LW1-LW3), and Palaung (PL) groups practice patrilocality, whereas the Htin (TN1- TN3), Karen (KSK1- KSK3, KPA, and KPW) and Lahu (MR and MB) are matrilocal. If genetic variation was influenced by residence pattern, then lower within-population genetic diversity coupled with greater genetic heterogeneity among populations is expected for matrilocal groups than for patrilocal groups for the mtDNA, whereas the opposite pattern is expected for the MSY (Oota et al. 2001). However, the MSY *h* and MPD values do not differ significantly between patrilocal and matrilocal groups (Table S6, Fig. S9). For mtDNA, genetic diversity values are significantly higher for patrilocal than for matrilocal groups for *h* but the differences are not statistically significant for MPD (Table S6). Notably, the patrilocal groups HM1, HM4 and LS exhibit higher than average MPD values for the MSY (78.65-99.37, compared to the average of 51.74) and some matrilocal groups, e.g. KSK3 and TN1-TN2, show much lower MPD (6.92-20.69) than average (33.99) for mtDNA (Table S1). Furthermore, the genetic diversity of all Htin groups are much lower than the other groups (Fig. S9).

For genetic differences between-populations revealed by AMOVA (Table 1), the MSY genetic variation among patrilocal populations is much higher than that for mtDNA (MSY: 27.43%, *P*<0.01; mtDNA: 6.36%, *P*<0.01), while for the matrilocal groups the mtDNA genetic variation is much higher, but still less than that for the MSY (MSY: 24.49%, *P*<0.01, mtDNA: 16.46%, *P*<0.01). Much stronger contrasting between-group variation is seen in two patrilocal groups, i.e. Hmong (MSY: 18.13%, *P*<0.01, mtDNA: 0.53%, *P*>0.05) and Lawa (MSY: 35.65%, *P*<0.01, mtDNA: 7.78%, *P*<0.01) and one matrilocal Htin group (MSY: 9.85%, *P*<0.01, mtDNA: 25.71%, *P*<0.01) (Fig. S10). In contrast to our previous study (Kutanan et al., 2019), with the inclusion of the new KSK3 population, the matrilocal Karen shows more differentiation for the MSY than for mtDNA (MSY: 9.11%, *P*<0.01, mtDNA: 5.85%, *P*<0.01). When Hmong, IuMien and Lahu from Thailand and Vietnam were combined, contrasting patterns of genetic variation between the MSY and mtDNA were still in accordance with expectations based on residence pattern, albeit the IuMien show only slightly higher between-group differentiation for the MSY than for mtDNA (Table 1).

Another potential effect of patrilocality vs. matrilocality is on the shared haplotypes between populations. If recent contact between populations is influenced by residence pattern, one would expect more MSY haplotype sharing among matrilocal groups than among patrilocal groups, and more mtDNA sharing among patrilocal groups than among matrilocal groups. However, the results for haplotype sharing between populations within matrilocal and patrilocal groups do not show a strong effect (Table S7). Haplotype sharing for the MSY is slightly lower on average for patrilocal groups (0.15) than for matrilocal groups (0.18), in accordance with expectations, but haplotype sharing for mtDNA is also lower on average for patrilocal groups (0.15) than for matrilocal groups (0.22), which is not in accordance with expectations based on residence pattern.

## Discussion

Our previous studies have focused on the genetic ancestry of the TK and AA groups in Thailand and Laos, here we investigate the less well-studied HM and ST speaking groups from Thailand, to gain more insights into the genetic history of MSEA. We sequenced ∼2.3 mB of the MSY and complete mtDNA genomes of the HM and ST groups who are regarded as hill tribes from northern Thailand, as well as additional TK groups from northern and northeastern Thailand. Although we focus on the HM and ST groups, we note that the previously-observed general pattern of overall genetic homogeneity of Thailand TK groups (Kutanan et al., 2019) continues to be maintained with these additional TK groups, consistent with the idea that the TK language family spread via demic diffusion (Kutanan et al., 2017). However, an additional insight arises when we compare the Thai TK data to similar mtDNA and MSY data from Vietnamese TK groups (Macholdt et al., 2019): some of the TK speaking groups from Vietnam (i.e. Nung, Tay and Thai) are quite similar to the TK populations from northeastern Thailand (Phutai, Kalueang and Black Tai) (Fig. 5). This is in agreement with historical evidence for a migration of the ancestors of the Phutai, Kalueang and Black Tai from Vietnam through Laos during last 200 years (Schliesinger, 2000; 2001).

### Genetic differences between the Hmong and IuMien groups and their origins

The Hmong-Mien (HM) language family is one of the major language families in MSEA, comprising some 39 languages (35 Hmongic and 4 Mienic) distributed across China, northern Vietnam, northern Laos, and northern Thailand (Ratliff, 2010). Although the Hmongic and Mienic languages are relatively similar to one another, there are differences due to divergent developments in the phonology (Ratliff, 2010). The heartland of the Hmong people is considered to be the southern Chinese province of Kweichow, where they were established at least 2000 years ago, probably arriving from the east (Schliesinger, 2000). Migrations into Thailand through Laos are documented since the second half of the 19^th^ century A.D. The IuMien are, like the Hmong, thought to have an origin in southeastern China, from which they started to migrate southwards to Vietnam in the 13^th^ century A.D., entering Thailand about 200 years ago (Schliesinger 2000; Besaggio et al., 2007).

Previous studies of HM groups have reported sequences of the mtDNA hypervariable region 1 with some diagnostic coding SNPs to define haplogroups (Wen et al., 2005), and Y-STR variation and genotypes for Y chromosomal bi-allelic loci (Cai et al., 2011). Here we analyze complete mtDNA and partial MSY sequences from five Hmong and two IuMien populations from Thailand; strikingly, we find significant differences between Hmong and Mien populations in Thailand, with the IuMien more similar to other populations, while the Hmong show genetic distinction that was not previously documented in Thai/Lao populations.

The genetic distances between the Hmong and one IuMien (Y2) are significantly different from zero for both mtDNA and the MSY, but the other IuMien population (Y1) are not significantly different from the Hmong. However, both IuMien populations also do not differ significantly from many other populations, suggesting contact with both HM and non-HM populations (Fig. 3B, Fig. 4). The two IuMien groups, but none of the Hmong groups, show significant negative Tajima’s D values for mtDNA (Fig. 2D); these may reflect population expansions due to contact. The haplogroup profiles and CA analyses also support a closer relationship between IuMien and non-HM speaking populations (Fig. S6 and S7; Table S5). The AMOVA results further indicate that the Hmong are closest to the ST in the paternal side, but closer to the AA and TK in the maternal side, whereas the IuMien are close to all other groups (Table 1).

Apart from the genetic distinction from their linguistic relatives, the IuMien, the Hmong in Thailand are genetically distinct from almost all other groups (Fig. 3). There are no shared mtDNA haplotypes between HM populations and other Thai/Lao populations, and only a few shared MSY haplotypes (Fig. 3A), Moreover, they do not overlap with other groups in the MDS analysis (Fig. 4), suggesting that they add unique genetic profiles that were not found in the previous studies of Thai/Lao AA, TK, and ST groups (Kutanan et al., 2019). This striking genetic divergence of Hmong populations in Thailand may reflect cultural isolation. Hmong communities have strong connections and prefer to marry with other Hmong groups rather than with non-Hmong groups (Geddes, 1976; Schliesinger, 2000). In contrast, the IuMien have shared haplotypes and closer genetic relatedness with several TK speaking groups, indicating more contact with other groups. These results may reflect the pronounced IuMien culture for adoption. Based on ethnographic accounts from the 1960s, around 10-20% of adult IuMien were adopted from other ethnic groups (both highland and lowland), in order to increase the size of their household and their family’s influence (Lewis, 1984; Schliesinger 2000; Jonsson, 2005; Besaggio et al., 2007). Another factor behind the genetic similarity of IuMien with other East Asian populations could be admixture, as suggested by sharing of features between IuMien (but not Hmong) and Sinitic languages (Blench, 2008).

Although the proto Hmong-Mien groups were suggested to have originated in central and southern China during the Neolithic Period (Wen et al., 2005), their greatest diversity is found between the Yangtze and Mekong rivers today. The ages of mtDNA haplogroups characteristic for HM groups, i.e. B4a5, B4g2, B5a1c1a*, B5a1c1a1, D4e1a3, and F1g1 are ∼6.63, ∼6.96, ∼10.67, ∼1.53, ∼6.83, and ∼11.54 kya, respectively (Fig. S3). The MSY haplogroups characteristic for HM groups, namely O2a2a1a2a1a2 (O-N5), O2a2a1a2a1a*, and C2e2 (C-F845), date to ∼2.45, ∼4.5, and ∼16.00 kya, respectively. However, if we focus on clades of haplogroup C-F845 that are exclusive to HM groups, the age is ∼2.85 kya (Fig. S2). It therefore seems that the HM predominant haplogroups originated during the Holocene to the late Neolithic Period, which is consistent with archaeological and historical evidence that the proto-HM group might be linked with the Neolithic cultures in the Middle Reach of the Yangtze River in southern China, namely the Daxi culture (5,300–6,400 YBP) and the Qujialing culture (4,600–5,000 YBP) (Wen et al., 2005). Our results are also consistent with a recent study of HM groups from Húnán, China (Xia et al., 2019), which identified lineages within MSY haplogroup O-N5 as specific to Hmong (and dated to ∼2.33 kya) and mtDNA haplogroup B5a1c1a as correlating with Pahng and IuMien (and dated to ∼9.80 kya). However, we do find B5a1c1a* and B5a1c1a1 specific to Hmong, but not IuMien, in Thailand. Overall, the coalescent ages of both MSY and mtDNA lineages are in the same range.

### The origin of the Sino-Tibetan groups

The ST family is both large (∼460 languages spoken by over a billion people) and spread across many countries in South, East and Southeast Asia, including China, Nepal, Bhutan, northeastern India, Pakistan, Myanmar, Bangladesh, Thailand, Vietnam, and Laos. There are two main ST subfamilies: Chinese and Tibeto-Burman, which separated around 6 kya based on lexical data (Wang, 1998). It has therefore been proposed that Neolithic people living at least 6 kya (Yang et al., 2012) or before 9 kya (van Driem, 2005) in northwestern China were probably the ancestors of modern ST populations. It has also been suggested that Sino-Tibetan languages originated among millet farmers, located in North China, around 7.2 kya (Sagart et al., 2019) or 5.9 kya (Zhang et al., 2019). Linguistic evidence then suggests differentiation between Sinitic and Tibeto- Burman languages, and also between Tibetan and Lolo-Burmese languages, and southward and westward expansions of ST groups (Sagart et al., 2019; Zhang et al., 2019).

Although an MSY lineage (O-M122*) was proposed to be characteristic of all modern ST populations (Su et al., 2000), subsequent studies have found further differentiation, e.g. haplogroup O2a1c-002611, which is at high frequency in Han Chinese but found at very low frequencies in Tibeto-Burman populations (Wang et al., 2013, 2014; Yan et al., 2011, 2014; Yao et al., 2017). Also, autosomal STR genotypes differentiate Tibetan and Lolo-Burmese speaking groups (Li et al., 2015; Yao et al., 2017).

ST-speaking groups in Thailand have not been studied in the same detail as those in China; here we analyzed three groups: Lisu, Lahu (Mussur) and Karen. Lisu and Lahu speak Lolo- Burmese languages, while the Karen languages belong to a different branch, Karenic (Schliesinger 2000). Historical evidence indicates that Lisu and Lahu migrated from southern China through Myanmar to northern Thailand about 100–200 years ago (Schliesinger, 2000). The Karen claim to be the first settlers in Myanmar who migrated from southern China before the arrival of Mon and Burmese people (Kuroiwa and Verkuyten, 2008), and the Karen groups in Thailand have been migrating from Myanmar started around 1750 A.D. due to the growing influence of the Burmese (Schliesinger, 2000).

The genetic distances between the Lisu and Lahu are significantly different from zero for both mtDNA and the MSY, and they also share both mtDNA and MSY haplotypes (Fig. 3A and 3B), indicating recent contact and/or shared ancestry. The two Lahu populations (Black Lahu (MB) and Red Lahu (MR)) are genetically similar to one another and both are genetically distinct from the other populations (Fig. 3B, Fig. 4B-4D). However, the Lisu do not differ significantly from many AA and TK populations (Fig. 3B), suggesting interactions between the Lisu and other populations. For the Karen, we have added an additional Karen population (Skaw (KSK3)) to the previously studied Karen populations; KSK3 has very low haplogroup diversity and MPD values for the MSY (Fig. 2), suggesting strong genetic drift that has in turn increased their divergence from the other Karen populations (Fig. 4). The other Karen groups are genetically similar to several populations, and also share many basal mtDNA M haplogroups (M21a, M* and M91a) with neighboring Austroasiatic populations, especially the Mon, suggesting significant admixing (Table S5). Previous studies based on autosomal STRs and SNPs also support the relatedness of the Karen and other AA groups in Thailand (Kutanan et al., 2015; Xu et al., 2010).

The estimated coalescent ages of the predominant lineages in ST populations provide an upper bound for their divergence/contact from other groups. MSY haplogroup O2a2b1a1-Page23, equivalent to O-M117, which was previously reported to be abundant in TB groups in southwestern China and in Han Chinese (Wang et al., 2014), was dated to around ∼2.41 kya in this study. However, this MSY lineage also occurs in HM, TK, and AA populations, reflecting recent shared ancestry and/or contact (Fig. S2). The ages of mtDNA haplogroups prevalent in Lahu and Lisu, namely A13, B4e, D4j1a1, and G1c, are dated to ∼6.74, ∼2.21, ∼2.49, and ∼2.20 kya, respectively (Fig. S3). Thus, the coalescent ages of many MSY and mtDNA lineages prevalent in ST groups are around the time of the Han expansion (∼2.5 kya; Bellwood, 2018).

### Contrasting paternal and maternal genetic variation in patrilocal vs. matrilocal groups

Previously, post-marital residence pattern has been shown to influence genetic variation in the hill tribes of Thailand (Oota et al., 2001; Besaggio et al., 2007; Kutanan et al., 2019). Our previous study had investigated four hill tribes: Lawa, Htin, Khmu ad Karen (Kutanan et al., 2019). Here we added data from four additional hill tribes (Hmong, IuMien, Lahu, and Lisu) for the most detailed investigation to date, comprising a total of 23 populations belonging to eight hill tribes. The Hmong, IuMien, Lisu, Lawa and Khmu are patrilocal (i.e., the wife moves to the residence of the husband after marriage) whereas the others are matrilocal. If postmarital residence pattern is having an influence on patterns of genetic variation, we would expect larger between-group differences and smaller within-group diversity for patrilocal groups for the MSY, and the same trends for matrilocal groups for mtDNA (Oota et al., 2001).

In general, the within-population genetic diversity values were not in agreement with expectations (Table S6; Fig. S10) whereas genetic differentiation between populations did go in the direction predicted by postmarital residence pattern for patrilocal groups but not for matrilocal groups (Table 1). However, when focusing on genetic differentiation within individual groups, the patrilocal Hmong and Lawa and the matrilocal Htin did fit with expectations, i.e. higher genetic differentiation among populations for the MSY than for mtDNA for the Hmong and Lawa, and the opposite for the Htin (Table 1). However, the matrilocal Karen show higher differentiation for the MSY than for mtDNA (Table 1), contrary to expectations and contrary to previous results based on four Karen populations (Kutanan et al., 2019). The addition of the KSK3 population increases the between-population MSY genetic variance from 2.3% to 9.1%, while the between-population mtDNA genetic variance is relatively unchanged. The low MSY MPD value (Fig. 2C) and outlier position in the MSY MDS plots (Fig. 4) indicate a strong effect of genetic drift on MSY variation in KSK3, which then mitigates any influence of postmarital residence pattern on MSY vs. mtDNA variation. Interestingly, overall we find less contrast between matrilocal and patrilocal groups than found previously for the hill tribes (Oota et al. 2001; Besaggio et al. 2007; Kutanan et al., 2019). Presumably this is because of both more detailed sampling and higher resolution analysis of the mtDNA and MSY genomes. And, this is not unexpected because while some studies find an impact of residence pattern on mtDNA/MSY variation, others do not (Kumar et al. 2006; Arias et al. 2018b). Many different factors can problably mtDNA/MSY variation, e.g. micro-evolutionarily factors such as genetic drift (as seen with the KSK3 population), physical landscape, subsistence strategies and other human cultural patterns (Wilkins and Marlowe 2006; Chaix et al. 2007) that diluted or erased any impact of residence pattern.

Nonetheless, one striking pattern remains in our data, and that concerns NEA vs. SEA ancestry. Previous genetic studies supported a north-south division in East Asian peoples and with some spread of northern ancestry to the south (Wen et al., 2004; 2005). Here we also find a higher frequency of both mtDNA and MSY lineages of SEA origin than of NEA origin in most of the studied populations. In general, the SEA specific maternal lineages (B5*, F1a*, M7b* and R9b*) are at an average frequency of 38.28%, while NEA mtDNA lineages (i.e. A*, D* and G*) have an average frequency of 9.38% (Table S5). The MSY haplogroups also show major SEA lineages (O1b*) predominating at an average frequency of 45.35%, and minor NEA lineages (C2e*, D- M174 and N*) at an average frequency of 8.33% (Table S3).

However, the HM and ST groups are a dramatic exception to this general pattern of higher SEA than NEA ancestry for both paternal and maternal lineages (Fig. 1). The estimated NEA maternal ancestry of the HM groups is 11.94%, comparable to that of other Thai/Lao populations (average = 9.11%), while the average frequency of NEA paternal lineages in HM groups is 24.72% (compared to the average frequency of 6.59% for other Thai/Lao populations). Conversely, in the ST groups we detect an average of 24.09% NEA maternal ancestry, which is much higher than the average NEA maternal ancestry for other Thai/Lao groups (7.57%), while the NEA paternal ancestry in ST groups is comparable to that in other Thai/Lao groups (11.95% vs. 7.88%).

Given that both HM and ST groups originated from southern China or northwestern China (Wen et al., 2004; Wen et al., 2005; Wang et al., 2014), we speculate that the ancestral HM and ST groups both had relatively high levels of NEA ancestry for both the MSY and mtDNA. However, due to subsequent contact with SEA groups, HM populations incorporated more SEA maternal than paternal lineages, while ST populations incorporated more SEA paternal than maternal lineages. This could be explained if the ancestral HM group was patrilocal (as HM populations are today), and so subsequent interactions between the HM ancestors and SEA groups incorporated more SEA mtDNA lineages than MSY lineages into HM populations. Conversely, if the ancestral ST group was matrilocal, subsequent interactions between ST ancestors and SEA groups would have incorporated more SEA MSY lineages than mtDNA lineages into ST populations. Matrilocality for ancient ST groups has also been suggested based on linguistic evidence (van Driem, 2007). The fact that some ST populations are now patrilocal (e.g., Lisu) while still exhibiting higher frequencies of NEA maternal lineages may then reflect recent changes from matrilocality to patrilocality.

## Conclusion

We have carried out the most extensive study to date, using high-resolution methods, of the maternal and paternal lineages in HM and ST speaking groups of northern Thailand. We find unexpected differences between the Hmong and IuMien, which may reflect different cultural practices, and genetic heterogeneity among ST groups. Compared to previous studies, we find less contrast in genetic diversity and differentiation between matrilocal and patrilocal groups among the hill tribes. However, a novel finding of this study is the contrast between HM and ST groups, both assumed to have origins in southern China, in frequencies of NEA maternal and paternal lineages. We suggest that this striking difference reflects ancestral patrilocality for HM groups vs. ancestral matrilocality for ST groups. Overall, our results further attest to the impact of cultural practices on patterns of mtDNA v. MSY variation in human populations.

## Materials and Methods

### Samples

Samples were collected from 416 individuals belonging to 14 populations classified into three linguistic groups: (1) Hmong-Mien groups, consisting of five Hmong populations (HM1-HM5) and two IuMien populations (Y1 and Y2); (2) Sino-Tibetan groups, consisting of two Lahu populations (MR and MB), one Lisu (LS) and one Karen subgroup Skaw (KPW3); and (3) Tai-Kadai groups, consisting of one Shan population (SH2), one Phutai population (PT2) and one Lao Isan population (IS5); (Table S1; Fig. 1). Genomic DNA samples of HM1-HM4 and Y1 and Y2 were from a previous study (Srikummool, 2005), while the remaining groups were newly-collected buccal samples obtained with written informed consent and with ethical approval from Khon Kaen University and the Ethics Commission of the University of Leipzig Medical Faculty. We extracted DNA using the Gentra Puregene Buccal Cell Kit (Qiagen, Germany) according to the manufacturer’s directions.

### Sequencing

Genomic libraries with double indices were prepared and enriched for mtDNA as described previously (Meyer and Kircher, 2010; Maricic et al., 2010). The libraries were sequenced on an Illumina Hiseq 2500 to obtain mtDNA consensus sequences as described by Arias et al. (2018a), with minor modifications as follows. We used Bustard for Illumina standard base calling and the read length was 76 bp. We then manually checked and manipulated sequences with Bioedit (www.mbio.ncsu.edu/BioEdit/bioedit.html). The sequence alignment to the Reconstructed Sapiens Reference Sequence (RSRS) (Behar et al., 2012) was done by MAFFT 7.271 (Katoh and Standley, 2013).

We enriched for ∼2.3 mB of the MSY from the same genomic libraries for male samples via in-solution hybridization-capture using a previously designed probe set (Kutanan et al. 2018a; 2019) and the Agilent Sure Select system (Agilent, CA). Sequencing was carried out on the Illumina HiSeq 2500 platform with paired-end reads of 125-bp length and we used Bustard for Illumina standard base calling. We used leeHOM to trim Illumina adapters and merge completely overlapping paired sequences (Renaud et al. 2014). We used deML to demultiplex the pooled sequencing data (Renaud et al. 2015). The alignment and post-processing pipeline of the sequencing data was carried out as previously described (Kutanan et al. 2018a).

### Statistical Analysis

The newly-generated 234 MSY sequences from 14 populations were combined with 928 sequences from 59 populations from our previous studies (Kutanan et al., 2018a; Kutanan et al., 2019) for a total of 1,162 sequences belonging to 73 populations. For mtDNA, combining the 416 new sequences from this study with 1,434 sequences from our previous studies (Kutanan et al., 2017; Kutanan et al., 2018a; Kutanan et al., 2018b) brings the total to 1,850 sequences from 73 populations. Summary statistics of the genetic diversity within populations, the matrix of pairwise genetic distances (*Φ*_st_), and analyses of molecular variance (AMOVA) were obtained with Arlequin 3.5.1.3 (Excoffier and Lischer, 2010). To visualize population relatedness, the R package (R Development Core Team 2016) was used to carry out nonparametric MDS analysis (based on the *Φ*_st_ distance matrices for the MSY and mtDNA) and to construct heat plots of the *Φ*_st_ distance matrix and the matrix of shared haplotypes. STATISTICA 13.0 (StatSoft, Inc., USA) was used to carry out a correspondence analysis based on MSY and mtDNA haplogroup frequencies. For the MSY, haplogroup assignment was performed by yHaplo (Poznik 2016) while mtDNA haplogroups were assigned by Haplogrep (Kloss-Brandstätter et al., 2011) and Haplofind (Vianello et al., 2013). To obtain a broader picture of population relationships within Southeast Asia, we included publicly-available sequences from relevant populations for the MSY (Karmin et al. 2015; Mallick et al. 2016; Poznik et al. 2016; Machold et al., 2019) and mtDNA (Zheng et al., 2011; Diroma et al., 2014; Summerer et al., 2014; Li et al., 2015; Machold et al., 2019). To construct Bayesian skyline plots (BSP) per population and maximum clade credibility (MCC) trees per haplogroup, based on Bayesian Markov Chain Monte Carlo (MCMC) analyses, we used BEAST 1.8.4 (Drummond et al. 2012). For BSP plots by population, we conducted analyses both pooling all populations within the same ethnicity (e.g., pooling HM1-HM5), and for the individual populations. BEAST input files were created with BEAUTi v1.8.2 after first running jModel test 2.1.7 in order to choose the most suitable model of sequence evolution (Darriba et al., 2012). For the MSY, we used an MSY mutation rate of 8.71 × 10^−10^ substitutions/bp/year (Helgason et al. 2015), and the BEAST input files were modified by an in-house script to add in the invariant sites found in our data set. Both strict and log normal relaxed clock models were run, with marginal likelihood estimation (Baele et al. 2012, 2013). After each BEAST run, the Bayes factor was computed from the log marginal likelihood of both models to choose the best-fitting BSP/MCC tree. For mtDNA, we executed BSP analyses per population and the BEAST runs by haplogroup, with mutation rates of 1.708 × 10^−8^ and 9.883 × 10^−8^ for data partitioned between the coding and noncoding regions, respectively (Soares et al., 2009). Tracer 1.6 was used to generate the BSP plot from the BEAST results. The Bayesian MCMC estimates (BE) and 95% highest posterior density (HPD) intervals of haplogroup coalescent times were calculated using the CongPy6 sequence (haplogroup A1b1-M14) (Karmin et al. 2015) for rooting the tree for MSY haplogroup C and the Mbuti-3 sequence (haplogorup B-M182) (Mallick et al. 2016) for rooting the tree for haplogroups F and O. The RSRS was used for rooting the tree for mtDNA. The Bayesian MCC trees were assembled with TreeAnnotator and drawn with FigTree v 1.4.3.

## Acknowledgements

We thank Roland Schröder, Enrico Macholdt and Leonardo Arias for technical assistance. This study was supported by the Max Planck Society. W.K. and R.S. were also supported by the Thailand Research Fund (RSA6180058 and RTA6080001). J.K. was supported by Chiang Mai University.

## Conflict of interest

The authors declare that they have no conflict of interest.

## Data Availability

All reads that aligned to the region of the MSY that was targeted by the capture-enrichment array were deposited in the European Nucleotide Archive (ENA) (study ID: XXXX). The mtDNA sequences were deposited in The National Center for Biotechnology Information (NCBI) (Accession number: XXXXX). The ENA and NCBI accession numbers will be provided during revision.

