## Supplementary Figure for "Cultural variation impacts paternal and maternal genetic lineages of the Hmong-Mien and Sino-Tibetan groups from Thailand"

### There are 10 Supplementary Figures and 7 Supplementary Tables in this manuscript.

### Supplementary Figures

**Figure S1** Bar plots of the distribution of major haplogroups for (A) MSYand (B) mtDNA. The new populations are placed at the left of the figure.

**Figure S2** Bayesian maximum clade credibility (MCC) tree of MSY sequences belonging to: (A) haplogroup C*; (B) haplogroup F; (C) haplogroup O2a2a*.

**Figure S3** Bayesian maximum clade credibility (MCC) trees of mtDNA sequences belonging to: (A) haplogroup A; (B) haplogroup B4; (C) haplogroup B5a1c1a; (D) haplogroup B6; (E) haplogroup C; (F) haplogroup D; (G) haplogroup F1g; (H) haplogroup G; and (I) haplogroup R9.

**Figure S4** The MDS plot for 73 Thai/Lao populations based on the MSY $\Phi_{st}$ distances for (A) dimension 1 vs. 2; (B) dimension 1 vs. 3; (C) dimension 2 vs. 3 and based on mtDNA $\Phi_{st}$ distances for (D) dimension 1 vs. 2; (E) dimension 1 vs. 3; (F) dimension 2 vs. 3.

**Figure S5** The heat plot of the five-dimensional MDS for the 73 Thai/Lao populations. (A) MSY; (B) mtDNA.

**Figure S6** Correspondence Analysis based on MSY haplogroup frequencies for (A) dimension 1 vs. 2; (B) dimension 1 vs. 3; (C) dimension 1 vs. 4; (D) dimension 2 vs. 3; (E) dimension 2 vs. 4; (F) dimension 3 vs. 4.

**Figure S7** Correspondence Analysis based on mtDNAhaplogroup frequencies for (A) dimension 1 vs. 2; (B) dimension 1 vs. 3; (C) dimension 1 vs. 4; (D) dimension 2 vs. 3; (E) dimension 2 vs. 4; (F) dimension 3 vs. 4.

**Figure S8** The BSPs based on the MSY and mtDNA for 14 populations. Solid lines are the median estimated effective population size (y axis) through time from the present in years (x axis). The 95% highest posterior density limits are indicated by light-purple lines.

**Figure S9** Bar plots of (A) haplotype diversity and (B) mean number of pairwise differences for patrilocal (blue) and matrilocal (orange) populations. Population abbreviations are in Table S1.

**Figure S10** Genetic variation among populations within groups, defined by ethnicity, language and cultural practice.

### Supplementary Tables

**Table S1** General information concerning the studied populations and genetic diversity values.

**Table S2** Sequencing coverage and MSY haplogroup of each individual.

**Table S3** MSY haplolgroup frequencies. The populations sequenced in this study are on the left side of the table, and newly-reported haplogroups are in red font.

**Table S4** Sequencing coverage and mtDNA haplogroup of each individual.

**Table S5** MtDNA haplolgroup frequencies. The populations sequenced in this study are on the left side of the table, and newly-reported haplogroups are in red font.

**Table S6** Mann-Whitney U test results for comparing genetic diversity values between different language families and residence patterns.

**Table S7** Haplotype sharing between populations of the same patrilocal or matrilocal group, for groups with at least two different populations sampled.

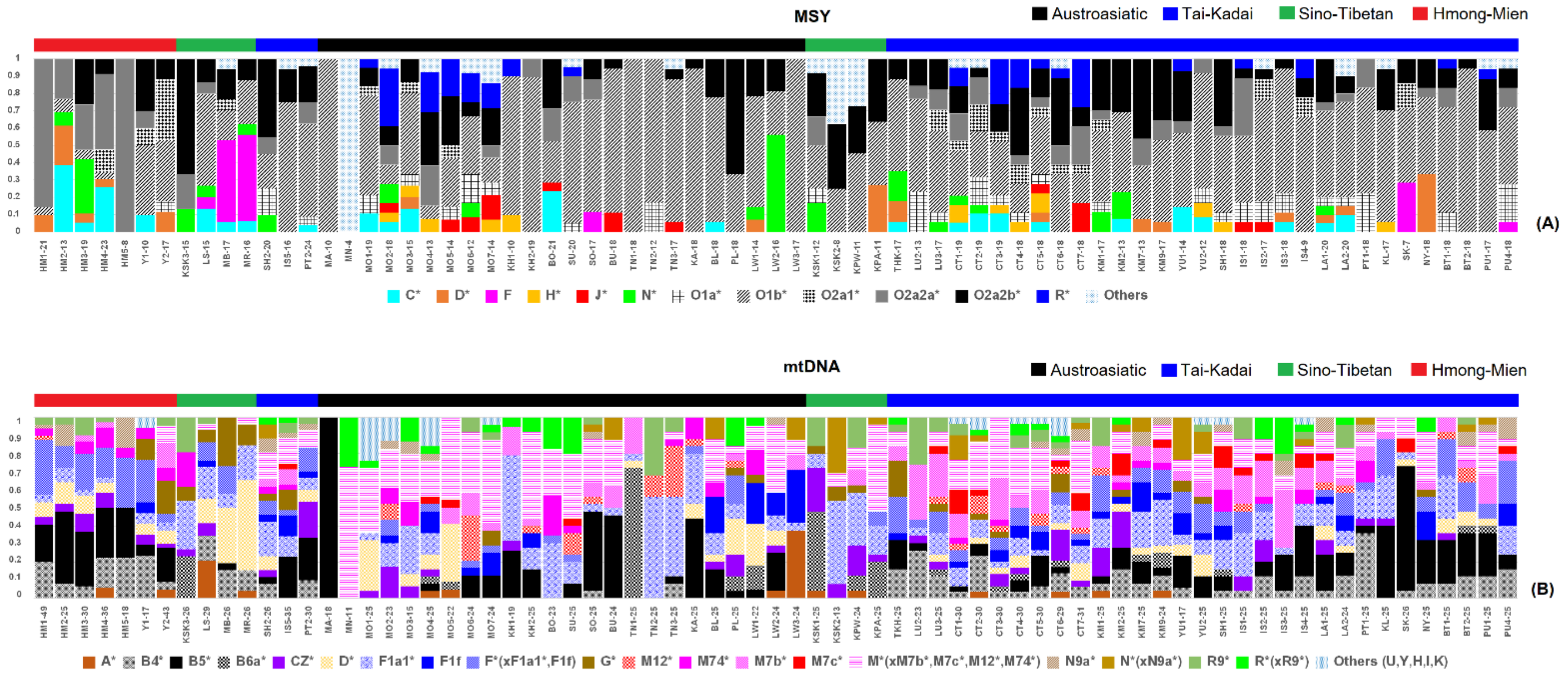


**Figure S1** Bar plots of the distribution of major haplogroups for (A) MSYand (B) mtDNA. The new populations are placed at the left of the figure.

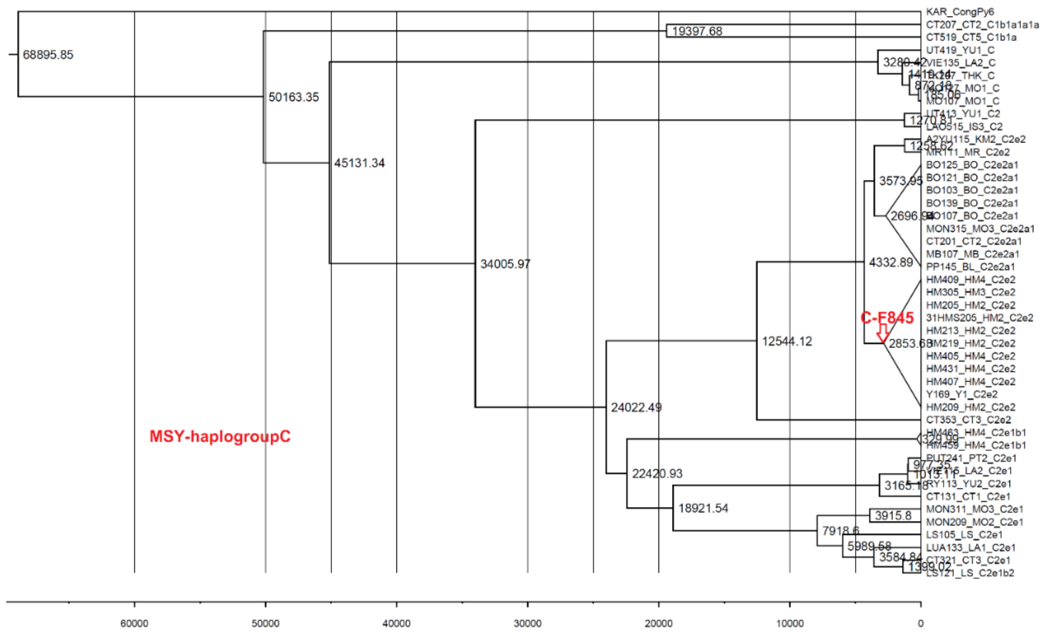
MSY-haplogroupC
C-F845
KAR_CongPy6
CT207_CT2_C1b1a1a1a
CT519_CT5_C1b1a
UT419_YU1_C
BO125_BO_C2e2a1
BO121_BO_C2e2a1
BO103_BO_C2e2a1
BO139_BO_C2e2a1
MON315_MO3_C2e2a1
CT201_CT2_C2e2a1
MB107_MB_C2e2a1
PP145_BL_C2e2a1
HM409_HM4_C2e2
HM305_HM3_C2e2
HM205_HM2_C2e2
31HMS205_HM2_C2e2
HM213_HM2_C2e2
HM405_HM4_C2e2
HM431_HM4_C2e2
HM407_HM4_C2e2
Y169_Y1_C2e2
HM209_HM2_C2e2
CT353_CT3_C2e2
CT131_CT1_C2e1
MON311_MO3_C2e1
MON209_MO2_C2e1
LS105_LS_C2e1
LUA133_LA1_C2e1
68895.85
50163.35
45131.34
34005.97
24022.49
22420.93
18921.54
12544.12
19397.68
4332.89
3573.95
3915.8
3165.18
60000
50000
40000
30000
20000
10000
0
(A)

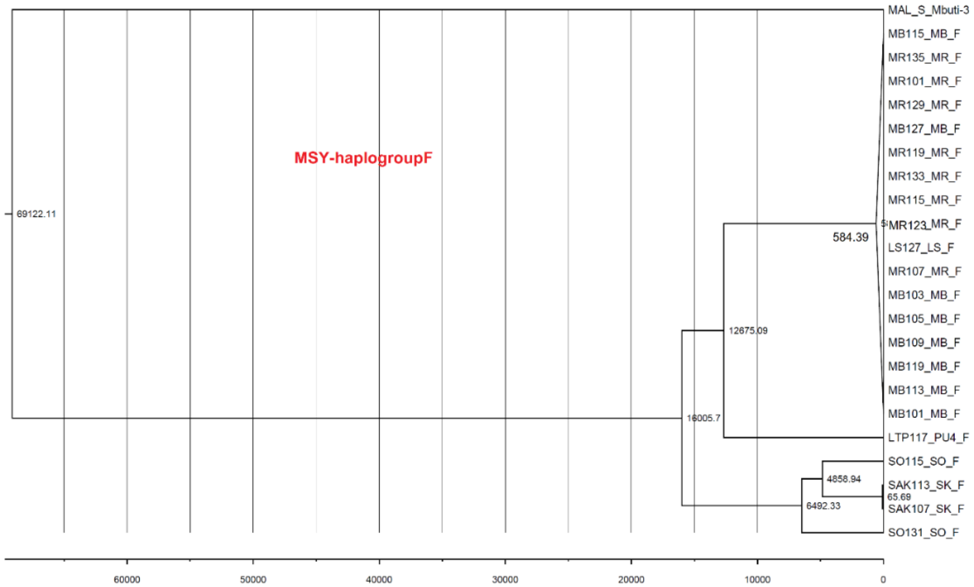
MSY-haplogroupF
MAL_S_Mbuti-3
MB115_MB_F
MR135_MR_F
MR101_MR_F
MR129_MR_F
MB127_MB_F
MR119_MR_F
MR133_MR_F
MR115_MR_F
MR123_MR_F
LS127_LS_F
MR107_MR_F
MB103_MB_F
MB105_MB_F
MB109_MB_F
MB119_MB_F
MB113_MB_F
MB101_MB_F
LTP117_PU4_F
SO115_SO_F
SAK113_SK_F
SAK107_SK_F
SO131_SO_F
69122.11
584.39
12675.09
16005.7
4858.94
65.69
6492.33
60000
50000
40000
30000
20000
10000
0
(B)

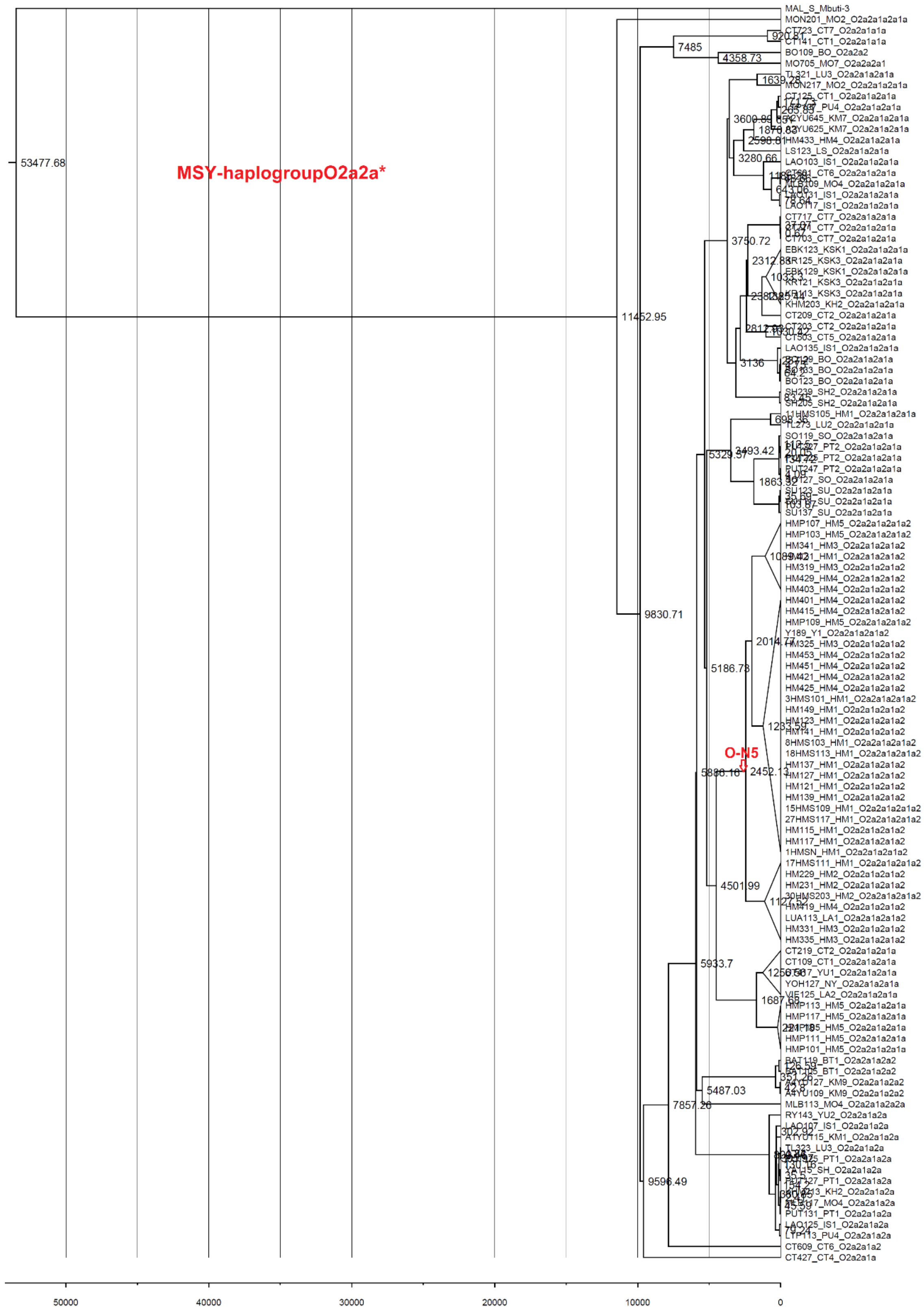


(C)

**Figure S2** Bayesian maximum clade credibility (MCC) tree of MSY sequences belonging to: (A) haplogroup C*; (B) haplogroup F; (C) haplogroup O2a2a*.

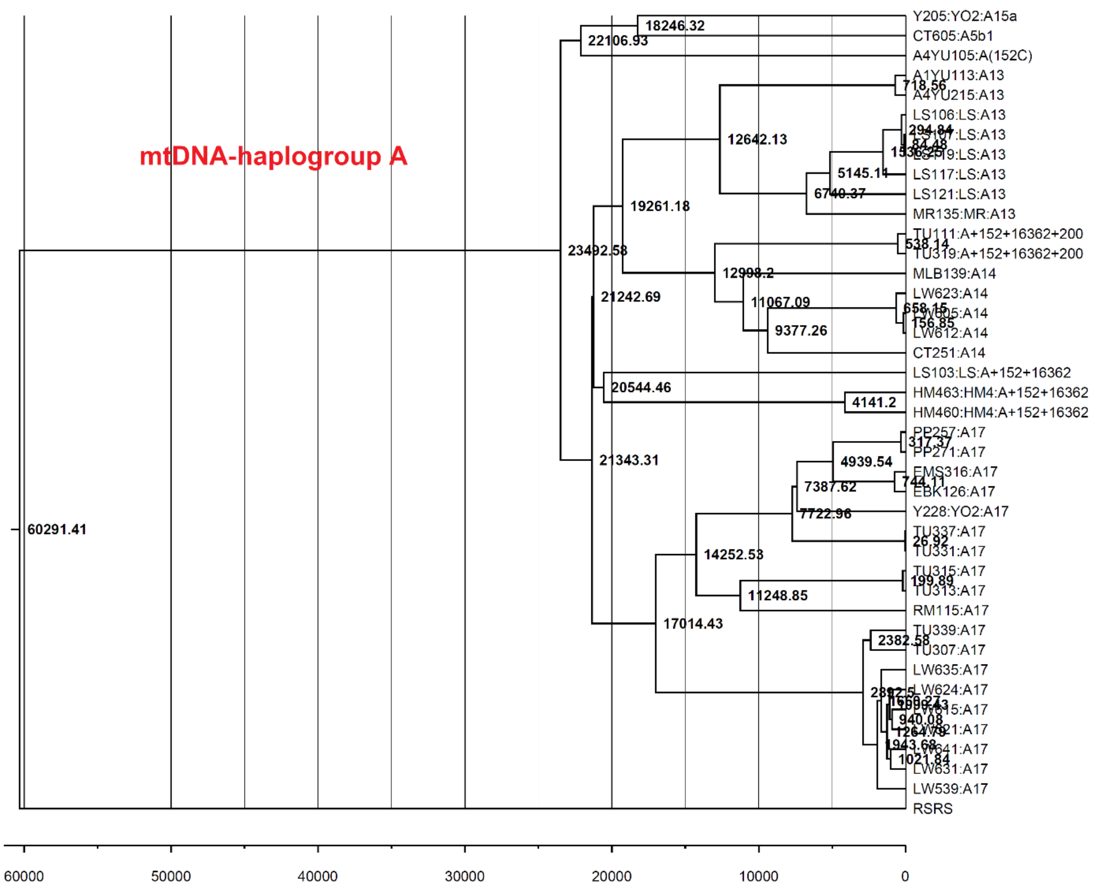
mtDNA-haplogroup A
Y205:YO2:A15a
CT605:A5b1
A4YU105:A(152C)
A1YU113:A13
A4YU215:A13
LS106:LS:A13
LS117:LS:A13
LS121:LS:A13
MR135:MR:A13
TU111:A+152+16362+200
TU319:A+152+16362+200
MLB139:A14
LW623:A14
LW612:A14
CT251:A14
LS103:LS:A+152+16362
HM463:HM4:A+152+16362
HM460:HM4:A+152+16362
PP257:A17
PP271:A17
EMS316:A17
EBK126:A17
Y228:YO2:A17
TU337:A17
TU331:A17
TU315:A17
TU313:A17
RM115:A17
TU339:A17
TU307:A17
LW635:A17
LW624:A17
LW631:A17
LW539:A17
RSRS
60291.41
23492.58
22106.93
18246.32
19261.18
12642.13
21242.69
12998.2
11067.09
9377.26
20544.46
4141.2
21343.31
17014.43
14252.53
11248.85
7387.62
7722.96
4939.54
5145.11
6740.37
2382.58
60000
50000
40000
30000
20000
10000
0

(A)

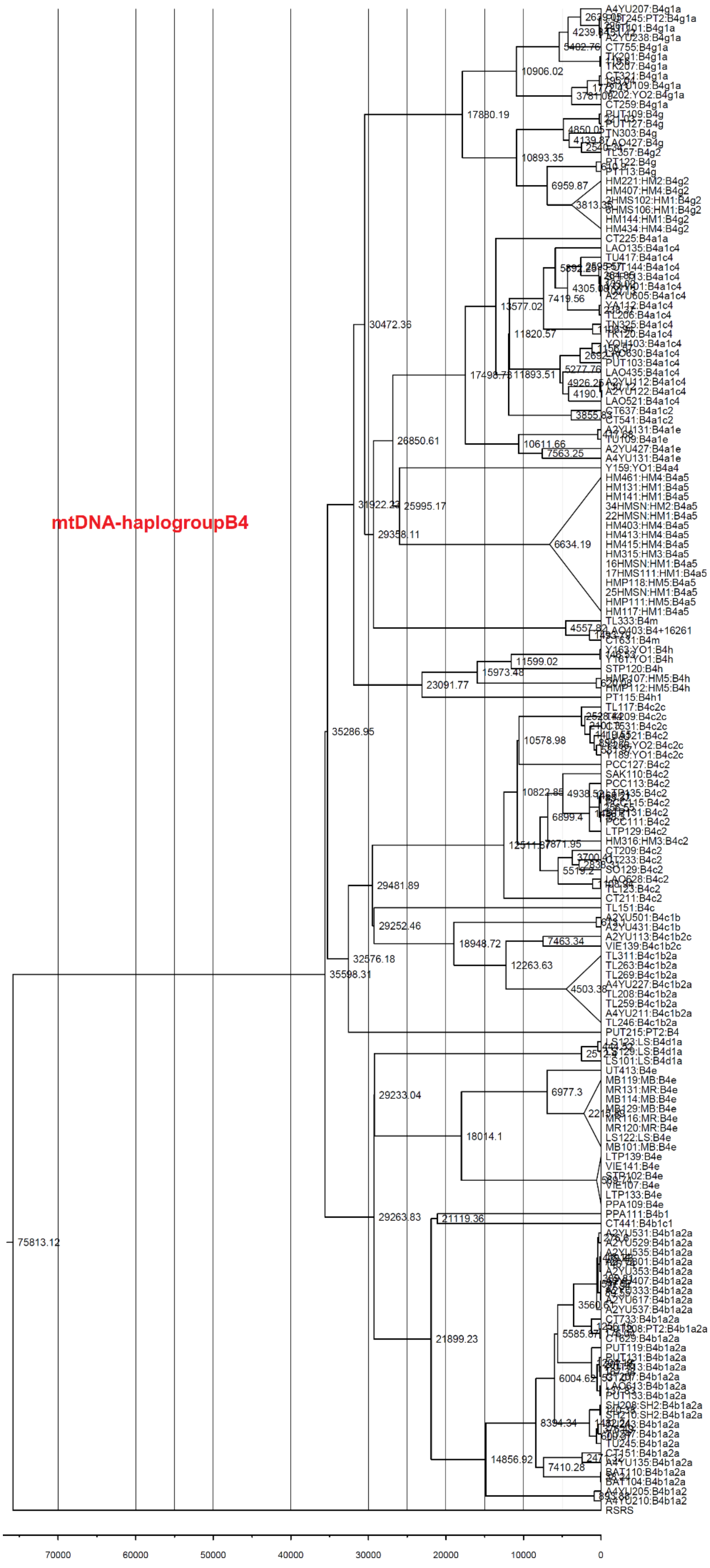

mtDNA-haplogroupB4
75813.12
35598.31
35286.95
32576.18
31922.22
30472.36
29358.11
28850.61
25995.17
17880.19
10906.02
10893.35
6959.87
5402.76
17498.78
13577.02
11820.57
11893.51
10611.66
7563.25
7419.56
5277.76
6634.19
23091.77
15973.48
11599.02
29481.89
29252.46
18948.72
12263.63
7463.34
4503.38
12511.87
10578.98
10822.85
6899.4
7871.95
5519.2
29233.04
18014.1
6977.3
29263.83
21119.36
21899.23
14856.92
8394.34
7410.28
6004.62
5585.87
A4YU207:B4g1a
A2YU238:B4g1a
CT755:B4g1a
TK201:B4g1a
TK207:B4g1a
CT321:B4g1a
CT259:B4g1a
TN303:B4g
TL357:B4g2
PT122:B4g
PT113:B4g
HM221:HM2:B4g2
HM407:HM4:B4g2
HM144:HM1:B4g2
HM434:HM4:B4g2
CT225:B4a1a
LAO135:B4a1c4
TU417:B4a1c4
A2YU605:B4a1c4
TL206:B4a1c4
TR120:B4a1c4
PUT103:B4a1c4
LAO435:B4a1c4
A2YU112:B4a1c4
A2YU122:B4a1c4
LAO521:B4a1c4
CT637:B4a1c2
CT541:B4a1c2
A2YU131:B4a1e
TU109:B4a1e
A2YU427:B4a1e
A4YU131:B4a1e
Y159:YO1:B4a4
HM461:HM4:B4a5
HM131:HM1:B4a5
HM141:HM1:B4a5
34HMSN:HM2:B4a5
22HMSN:HM1:B4a5
HM403:HM4:B4a5
HM413:HM4:B4a5
HM415:HM4:B4a5
HM315:HM3:B4a5
16HMSN:HM1:B4a5
17HMS111:HM1:B4a5
HMP118:HM5:B4a5
25HMSN:HM1:B4a5
HMP111:HM5:B4a5
HM117:HM1:B4a5
TL333:B4m
AO403:B4+16261
CT631:B4m
Y163:YO1:B4h
Y161:YO1:B4h
STP120:B4h
HMP112:HM5:B4h
PT115:B4h1
TL117:B4c2c
Y189:YO1:B4c2c
PCC127:B4c2
SAK110:B4c2
PCC113:B4c2
PCC111:B4c2
LTP129:B4c2
HM316:HM3:B4c2
CT209:B4c2
SO129:B4c2
LAO628:B4c2
TL123:B4c2
CT211:B4c2
TL151:B4c
A2YU501:B4c1b
A2YU431:B4c1b
A2YU113:B4c1b2c
VIE139:B4c1b2c
TL311:B4c1b2a
TL263:B4c1b2a
TL269:B4c1b2a
A4YU227:B4c1b2a
TL208:B4c1b2a
TL259:B4c1b2a
A4YU211:B4c1b2a
TL246:B4c1b2a
PUT215:PT2:B4
LS123:LS:B4d1a
LS101:LS:B4d1a
UT413:B4e
MB119:MB:B4e
MR131:MR:B4e
MB114:MB:B4e
MB129:MB:B4e
MR116:MR:B4e
MR120:MR:B4e
LS122:LS:B4e
MB101:MB:B4e
LTP139:B4e
VIE141:B4e
VIE107:B4e
LTP133:B4e
PPA109:B4e
PPA111:B4b1
CT441:B4b1c1
A2YU531:B4b1a2a
A2YU529:B4b1a2a
A2YU535:B4b1a2a
A2YU353:B4b1a2a
A2YU617:B4b1a2a
A2YU537:B4b1a2a
CT733:B4b1a2a
CT629:B4b1a2a
PUT119:B4b1a2a
PUT131:B4b1a2a
LAO613:B4b1a2a
PUT133:B4b1a2a
SH208:SH2:B4b1a2a
SH210:SH2:B4b1a2a
TU245:B4b1a2a
CT151:B4b1a2a
A4YU135:B4b1a2a
BAT110:B4b1a2a
BAT104:B4b1a2a
A4YU205:B4b1a2
A4YU210:B4b1a2
RSRS
70000
60000
50000
40000
30000
20000
10000
0


(B)

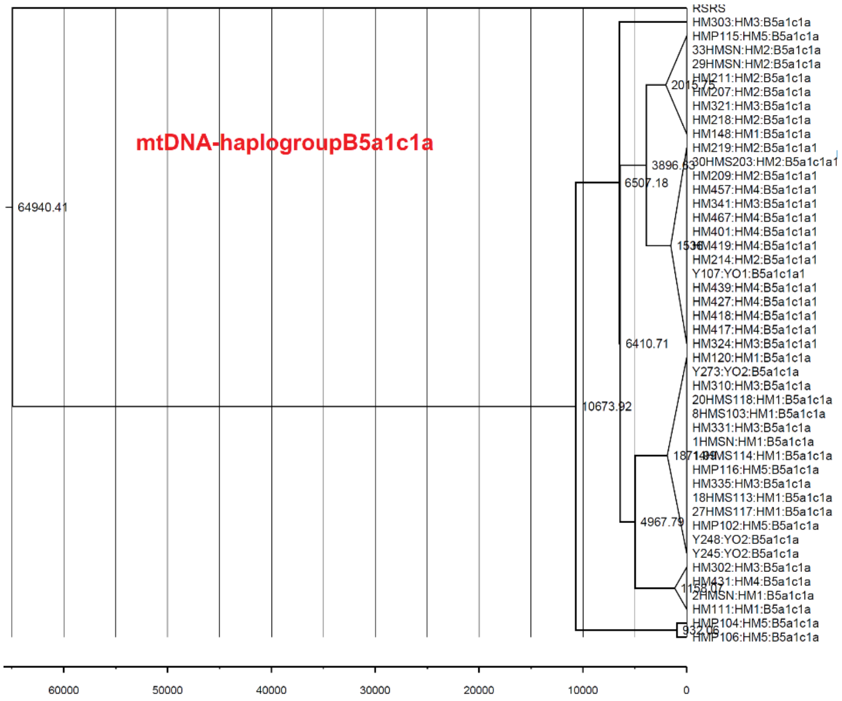
mtDNA-haplogroupB5a1c1a
RSRS
HM303:HM3:B5a1c1a
HMP115:HM5:B5a1c1a
33HMSN:HM2:B5a1c1a
29HMSN:HM2:B5a1c1a
HM207:HM2:B5a1c1a
HM321:HM3:B5a1c1a
HM218:HM2:B5a1c1a
HM148:HM1:B5a1c1a
HM219:HM2:B5a1c1a1
HM209:HM2:B5a1c1a1
HM457:HM4:B5a1c1a1
HM341:HM3:B5a1c1a1
HM467:HM4:B5a1c1a1
HM401:HM4:B5a1c1a1
HM214:HM2:B5a1c1a1
Y107:YO1:B5a1c1a1
HM439:HM4:B5a1c1a1
HM427:HM4:B5a1c1a1
HM418:HM4:B5a1c1a1
HM417:HM4:B5a1c1a1
HM324:HM3:B5a1c1a1
HM120:HM1:B5a1c1a
Y273:YO2:B5a1c1a
HM310:HM3:B5a1c1a
20HMS118:HM1:B5a1c1a
8HMS103:HM1:B5a1c1a
HM331:HM3:B5a1c1a
1HMSN:HM1:B5a1c1a
HMP116:HM5:B5a1c1a
HM335:HM3:B5a1c1a
18HMS113:HM1:B5a1c1a
27HMS117:HM1:B5a1c1a
HMP102:HM5:B5a1c1a
Y248:YO2:B5a1c1a
Y245:YO2:B5a1c1a
HM302:HM3:B5a1c1a
HM111:HM1:B5a1c1a
64940.41
10673.92
6507.18
6410.71
4967.79
60000
50000
40000
30000
20000
10000
0

(C)

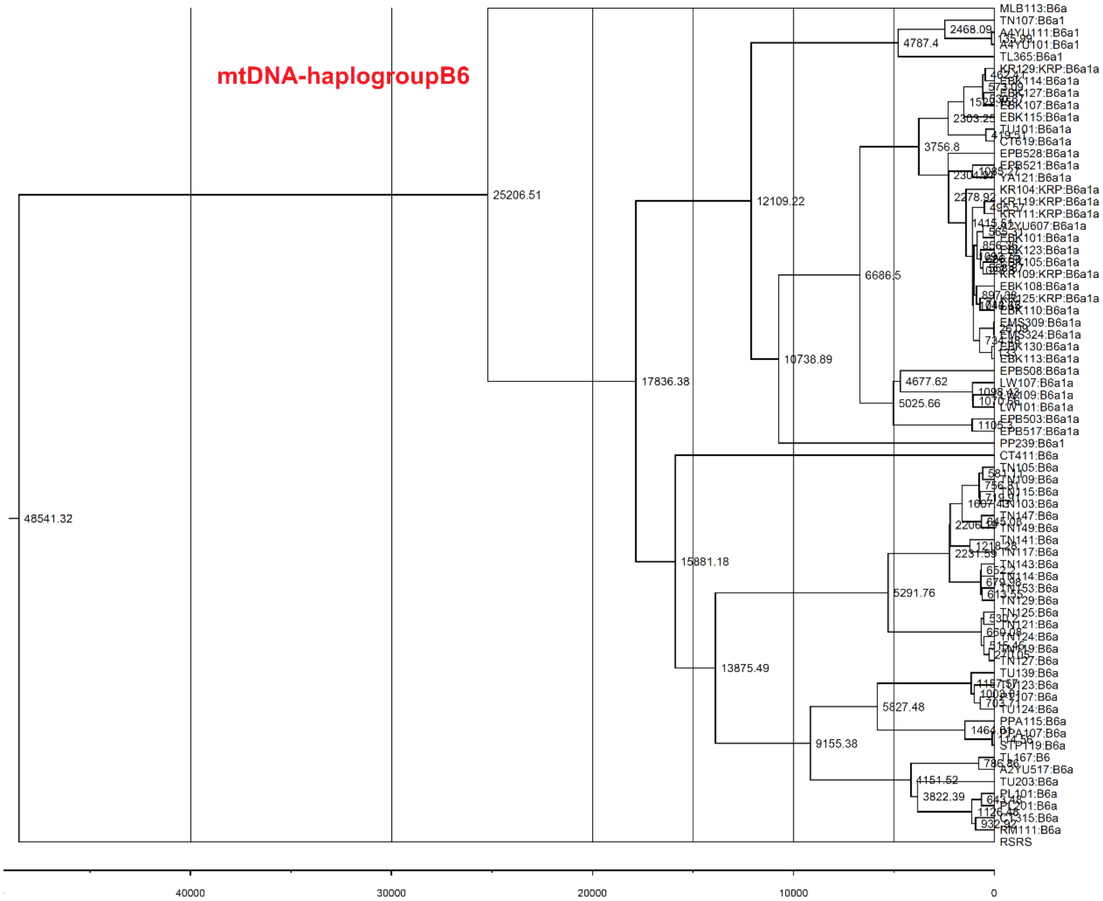
mtDNA-haplogroupB6
MLB113:B6a
TN107:B6a1
TL365:B6a1
EBK115:B6a1a
EPB528:B6a1a
KR104:KRP:B6a1a
EBK108:B6a1a
EBK113:B6a1a
EPB508:B6a1a
LW107:B6a1a
PP239:B6a1
CT411:B6a
TN129:B6a
TN127:B6a
TU139:B6a
TU124:B6a
PPA115:B6a
TU203:B6a
RSRS
48541.32
25206.51
17836.38
15881.18
13875.49
12109.22
10738.89
9155.38
6686.5
5827.48
5291.76
5025.66
4787.4
4677.62
4151.52
3822.39
3756.8
2468.09
40000
30000
20000
10000
0

(D)

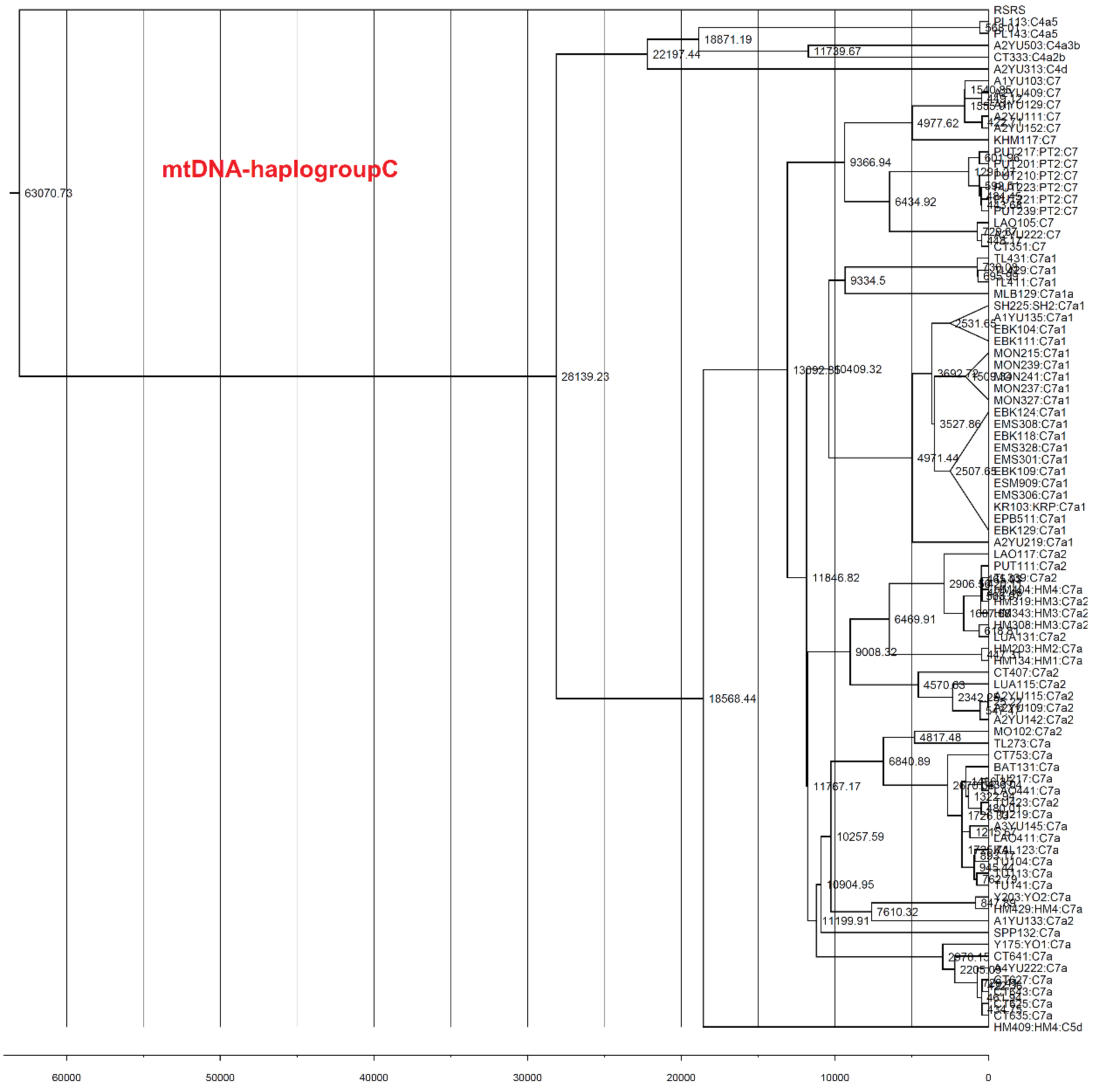
mtDNA-haplogroupC
63070.73
28139.23
22197.44
18871.19
11739.67
18568.44
13092.85
10409.32
9366.94
4977.62
6434.92
9334.5
4971.44
3692.72
3527.86
2531.65
2507.65
11846.82
6469.91
9008.32
2906.56
4570.63
2342.85
11767.17
6840.89
4817.48
10257.59
10904.95
7610.32
11199.91
2205.09
RSRS
PL113:C4a5
PL143:C4a5
A2YU503:C4a3b
CT333:C4a2b
A2YU313:C4d
A1YU103:C7
A2YU409:C7
A2YU111:C7
A2YU152:C7
KHM117:C7
PUT217:PT2:C7
PUT201:PT2:C7
PUT210:PT2:C7
PUT239:PT2:C7
LAO105:C7
CT351:C7
TL431:C7a1
TL411:C7a1
MLB129:C7a1a
SH225:SH2:C7a1
A1YU135:C7a1
EBK104:C7a1
EBK111:C7a1
MON215:C7a1
MON239:C7a1
MON237:C7a1
MON327:C7a1
EBK124:C7a1
EMS308:C7a1
EBK118:C7a1
EMS328:C7a1
EMS301:C7a1
EBK109:C7a1
ESM909:C7a1
EMS306:C7a1
KR103:KRP:C7a1
EPB511:C7a1
EBK129:C7a1
A2YU219:C7a1
LAO117:C7a2
PUT111:C7a2
HM319:HM3:C7a2
HM343:HM3:C7a2
HM308:HM3:C7a2
LUA131:C7a2
HM203:HM2:C7a
HM134:HM1:C7a
CT407:C7a2
LUA115:C7a2
A2YU115:C7a2
A2YU142:C7a2
MO102:C7a2
TL273:C7a
CT753:C7a
BAT131:C7a
A3YU145:C7a
LAO411:C7a
Y203:YO2:C7a
HM429:HM4:C7a
A1YU133:C7a2
SPP132:C7a
Y175:YO1:C7a
CT641:C7a
A4YU222:C7a
CT635:C7a
HM409:HM4:C5d
60000
50000
40000
30000
20000
10000
0

(E)

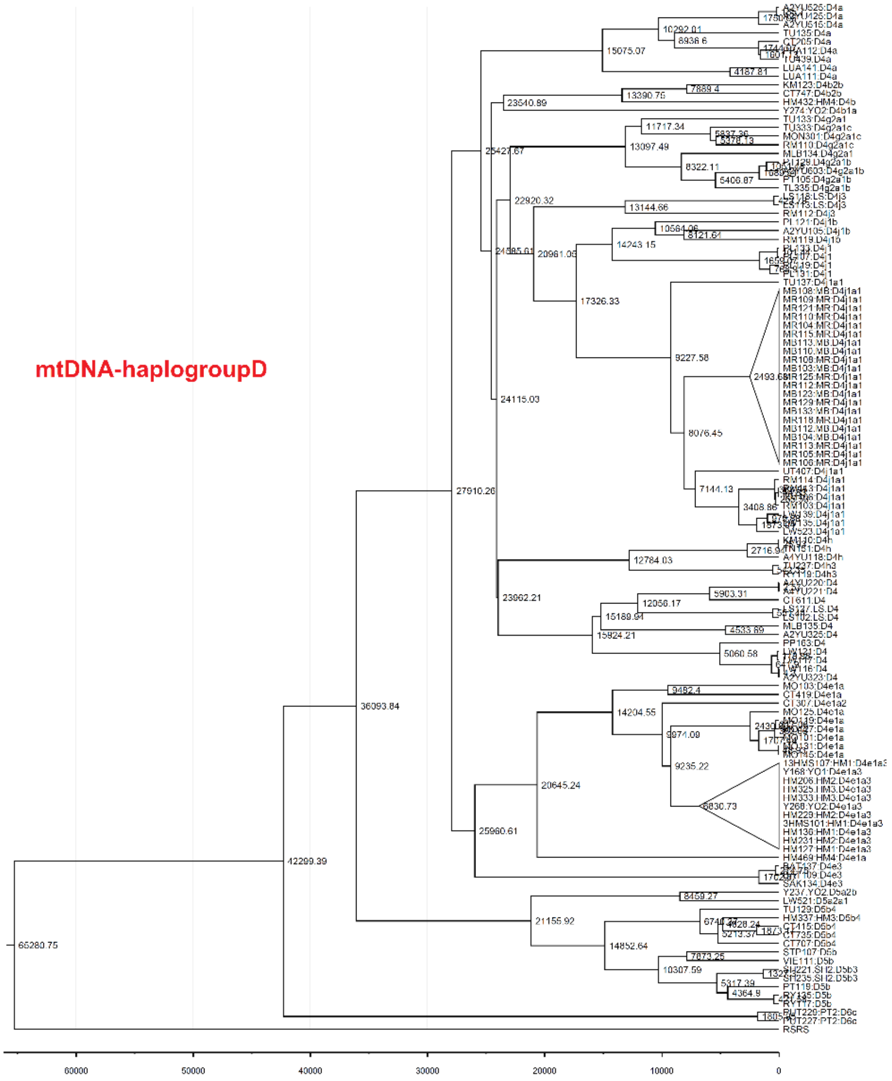
mtDNA-haplogroupD
60000
50000
40000
30000
20000
10000
0

(F)

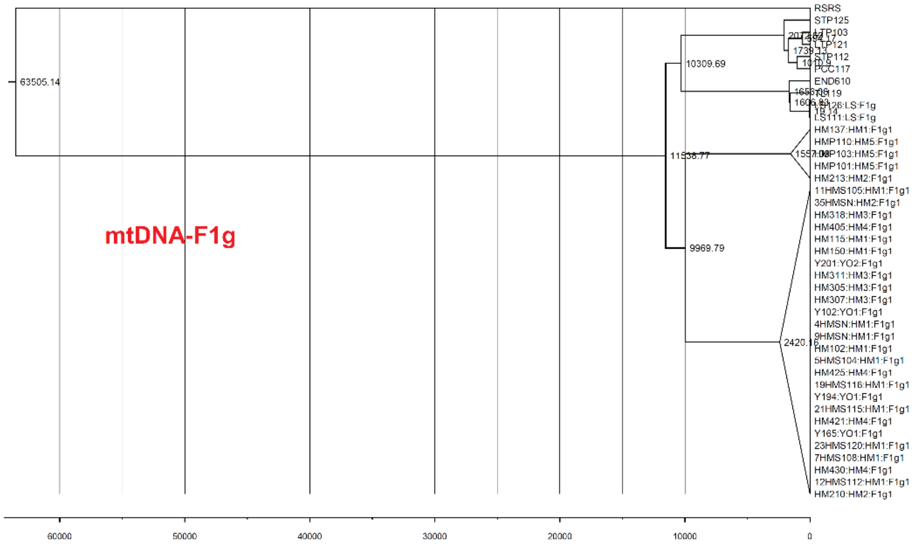
mtDNA-F1g
60000
50000
40000
30000
20000
10000
0

(G)

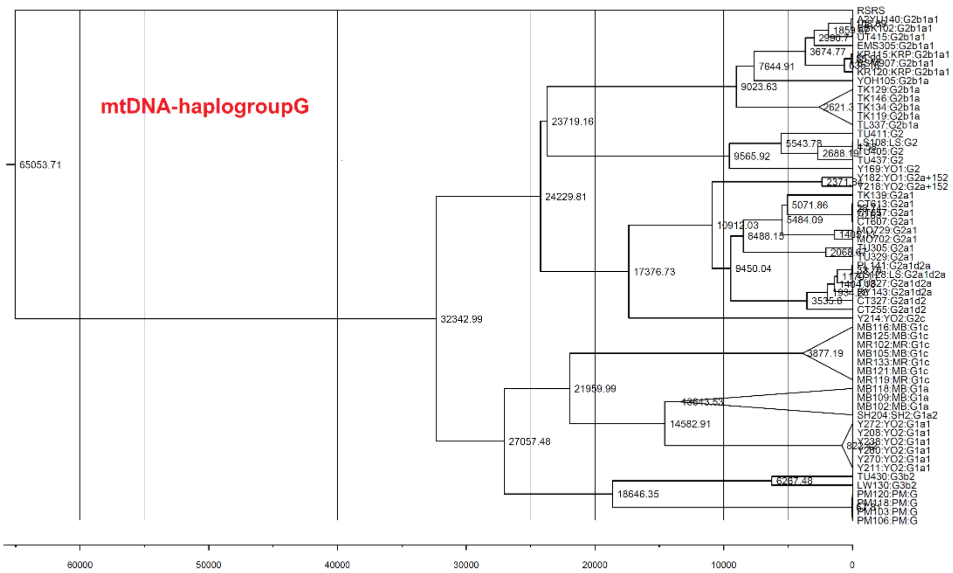


(H)

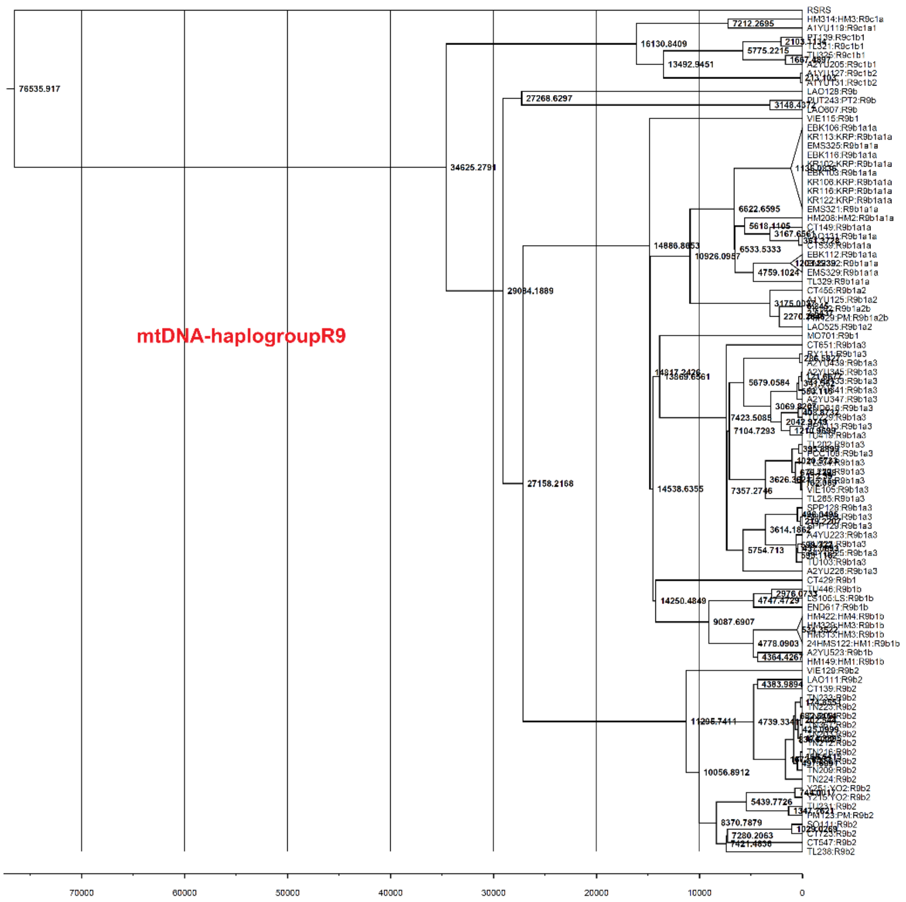


(I)

**Figure S3** Bayesian maximum clade credibility (MCC) trees of mtDNA sequences belonging to: (A) haplogroup A; (B) haplogroup B4; (C) haplogroup B5a1c1a; (D) haplogroup B6; (E) haplogroup C; (F) haplogroup D; (G) haplogroup F1g; (H) haplogroup G; and (I) haplogroup R9.

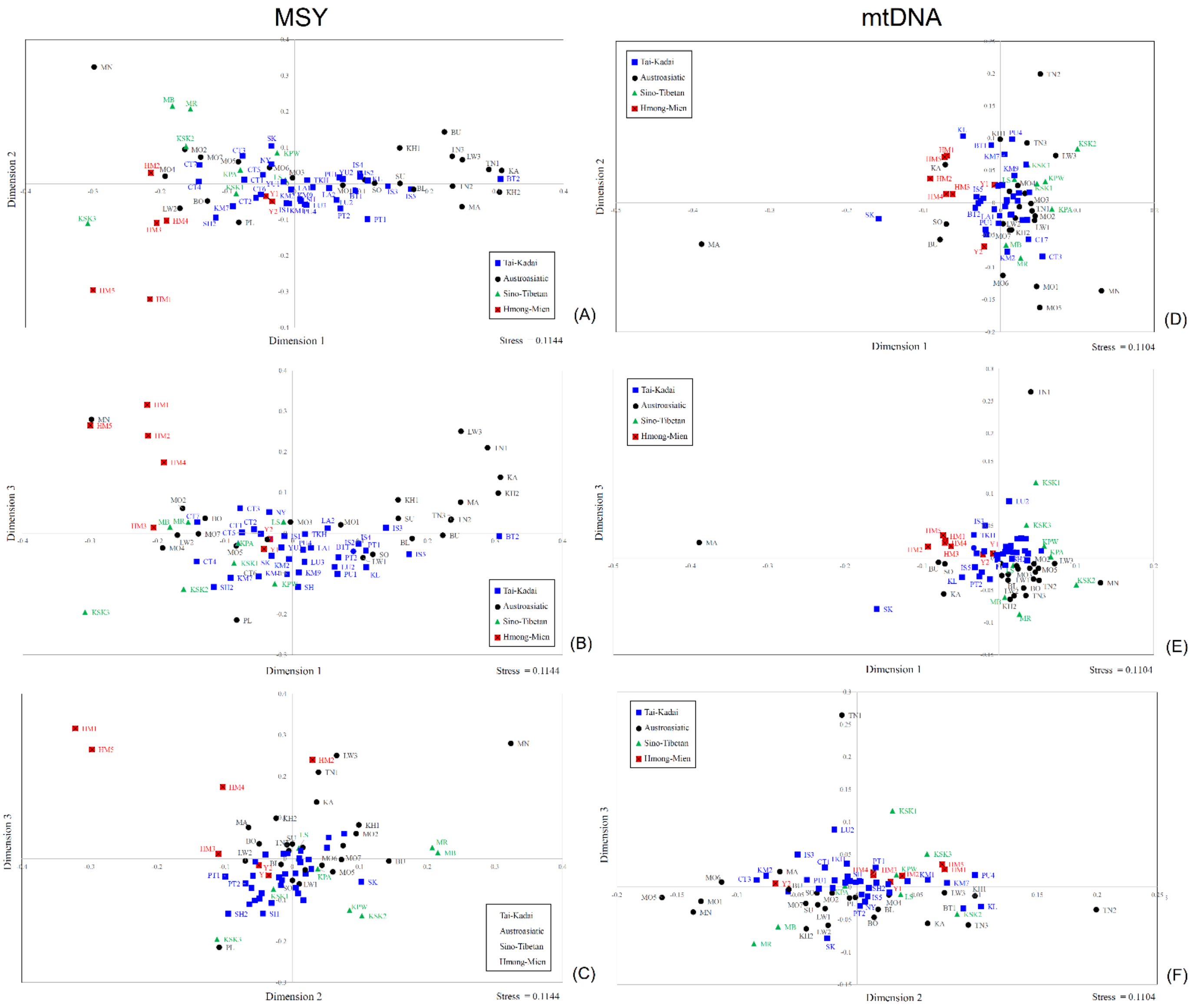


**Figure S4** The MDS plot for 73 Thai/Lao populations based on the MSY $\Phi_{st}$ distances for (A) dimension 1 vs. 2; (B) dimension 1 vs. 3; (C) dimension 2 vs. 3 and based on mtDNA $\Phi_{st}$ distances for (D) dimension 1 vs. 2; (E) dimension 1 vs. 3; (F) dimension 2 vs. 3.

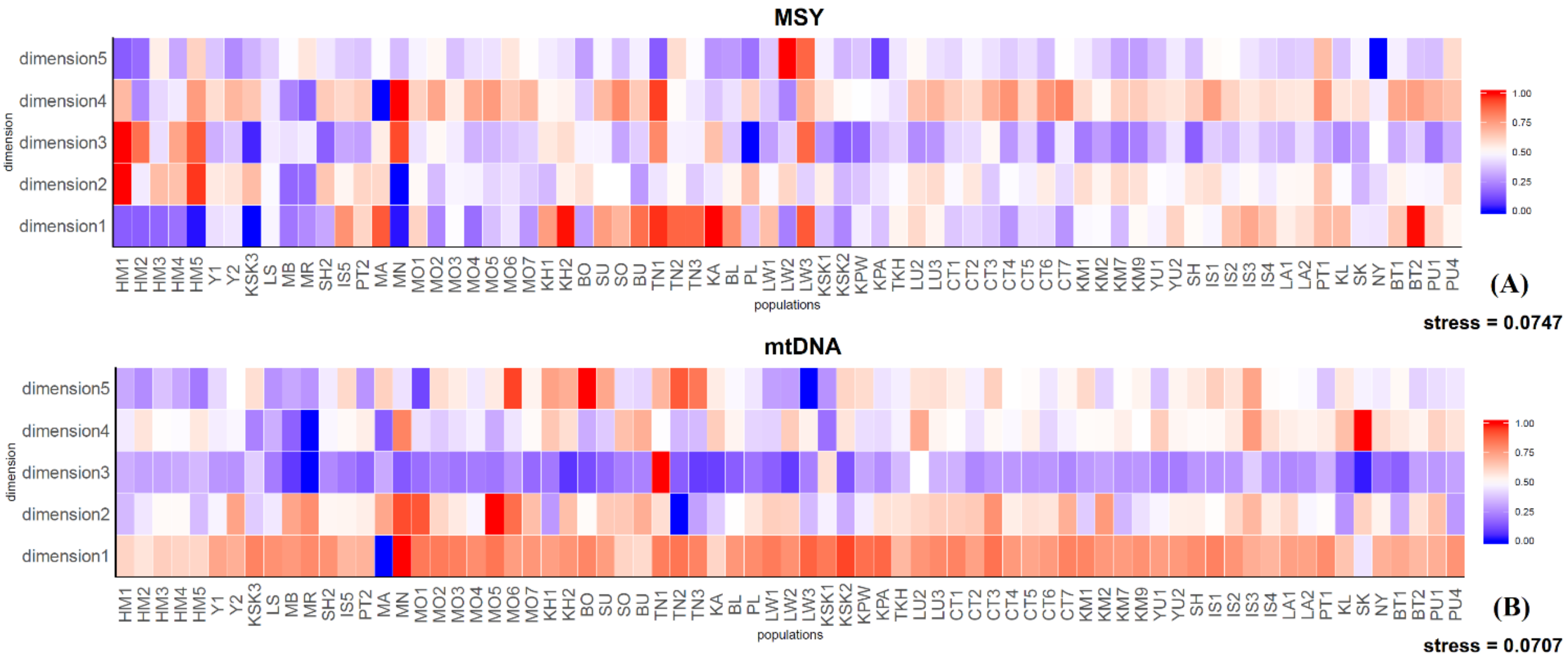


**Figure S5** The heat plot of the five-dimensional MDS for the 73 Thai/Lao populations. (A) MSY; (B) mtDNA.

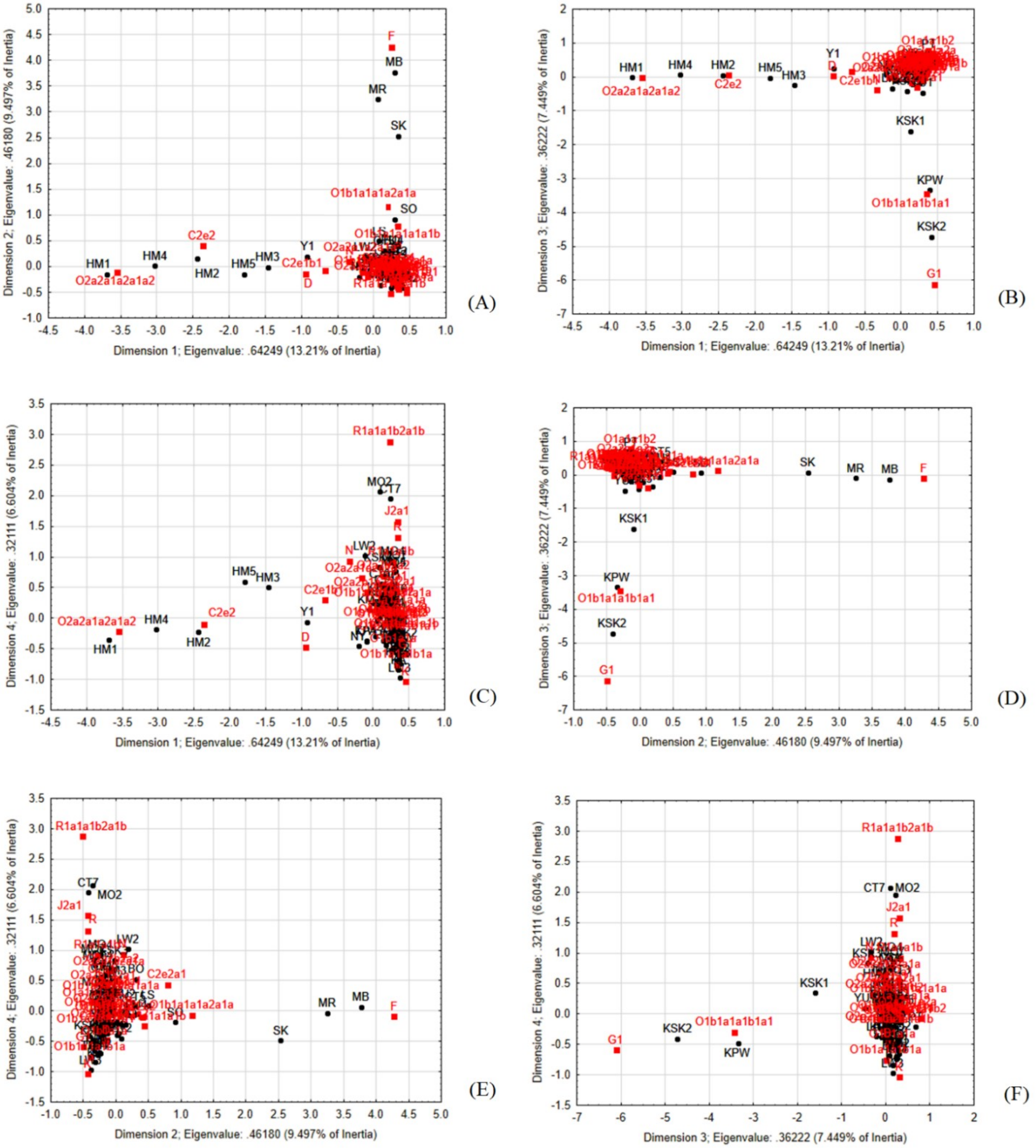


**Figure S6** Correspondence Analysis based on MSY haplogroup frequencies for (A) dimension 1 vs. 2; (B) dimension 1 vs. 3; (C) dimension 1 vs. 4; (D) dimension 2 vs. 3; (E) dimension 2 vs. 4; (F) dimension 3 vs. 4.

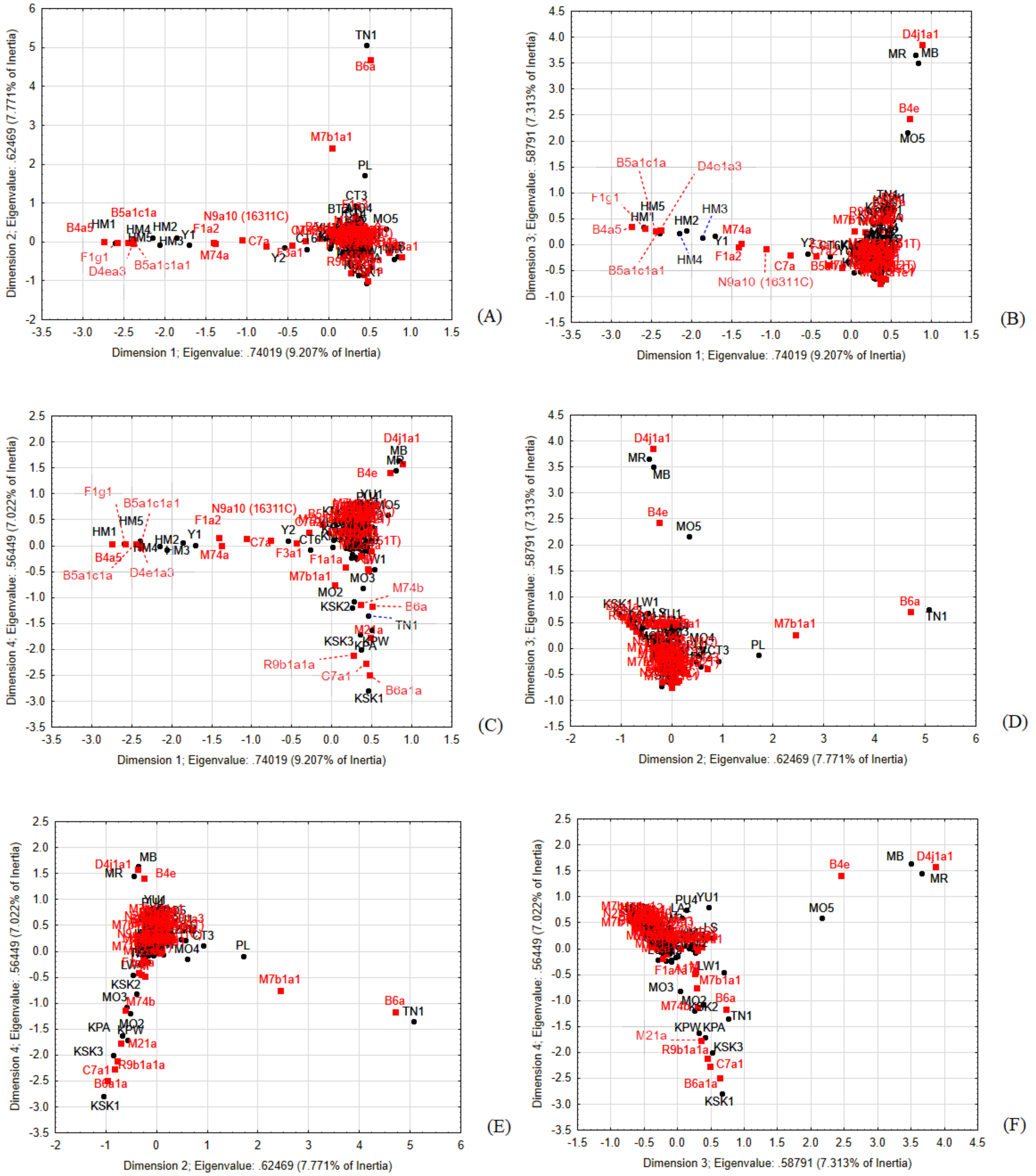


**Figure S7** Correspondence Analysis based on mtDNAhaplogroup frequencies for (A) dimension 1 vs. 2; (B) dimension 1 vs. 3; (C) dimension 1 vs. 4; (D) dimension 2 vs. 3; (E) dimension 2 vs. 4; (F) dimension 3 vs. 4.

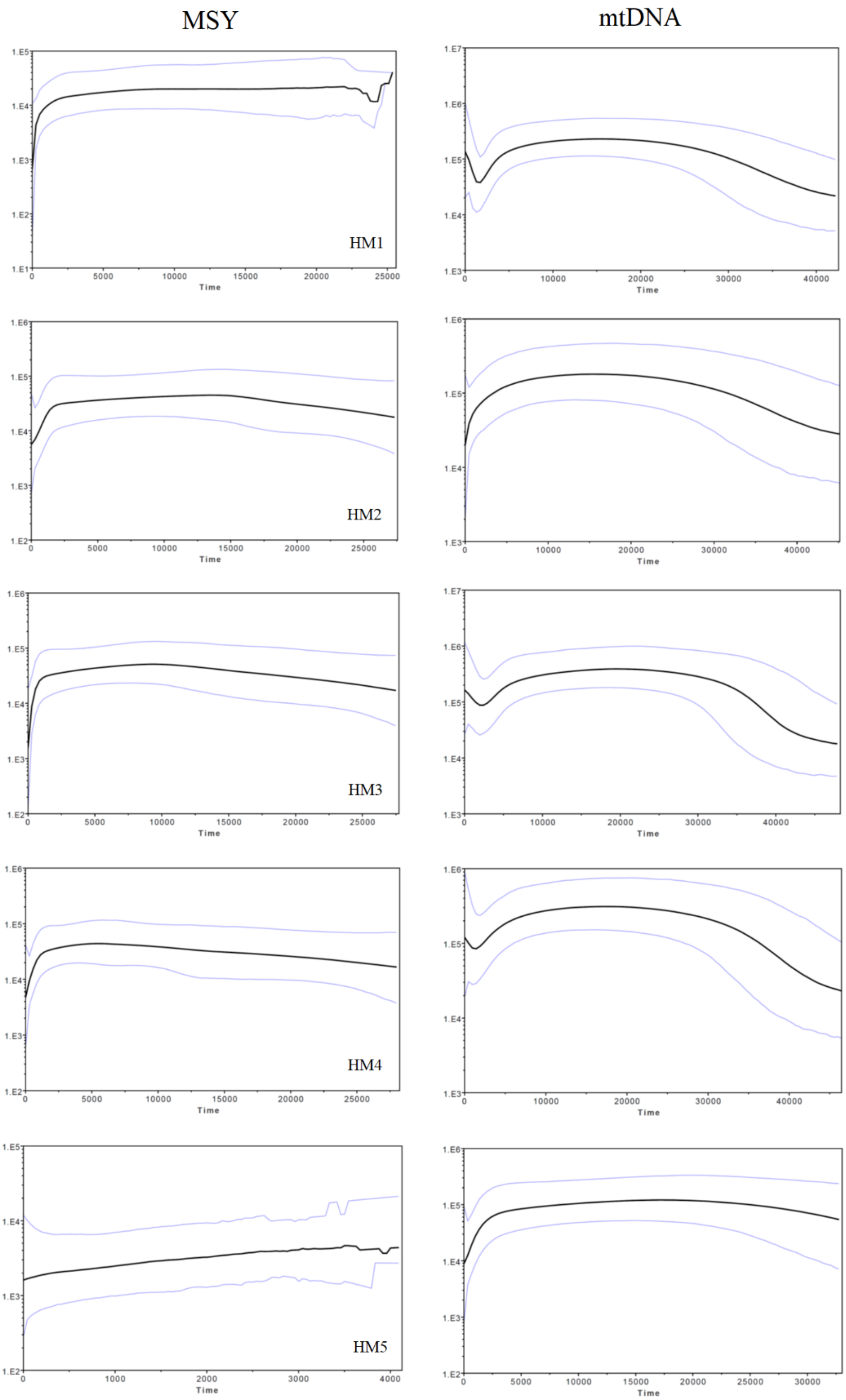
MSY
mtDNA
HM1
HM2
HM3
HM4
HM5
Time

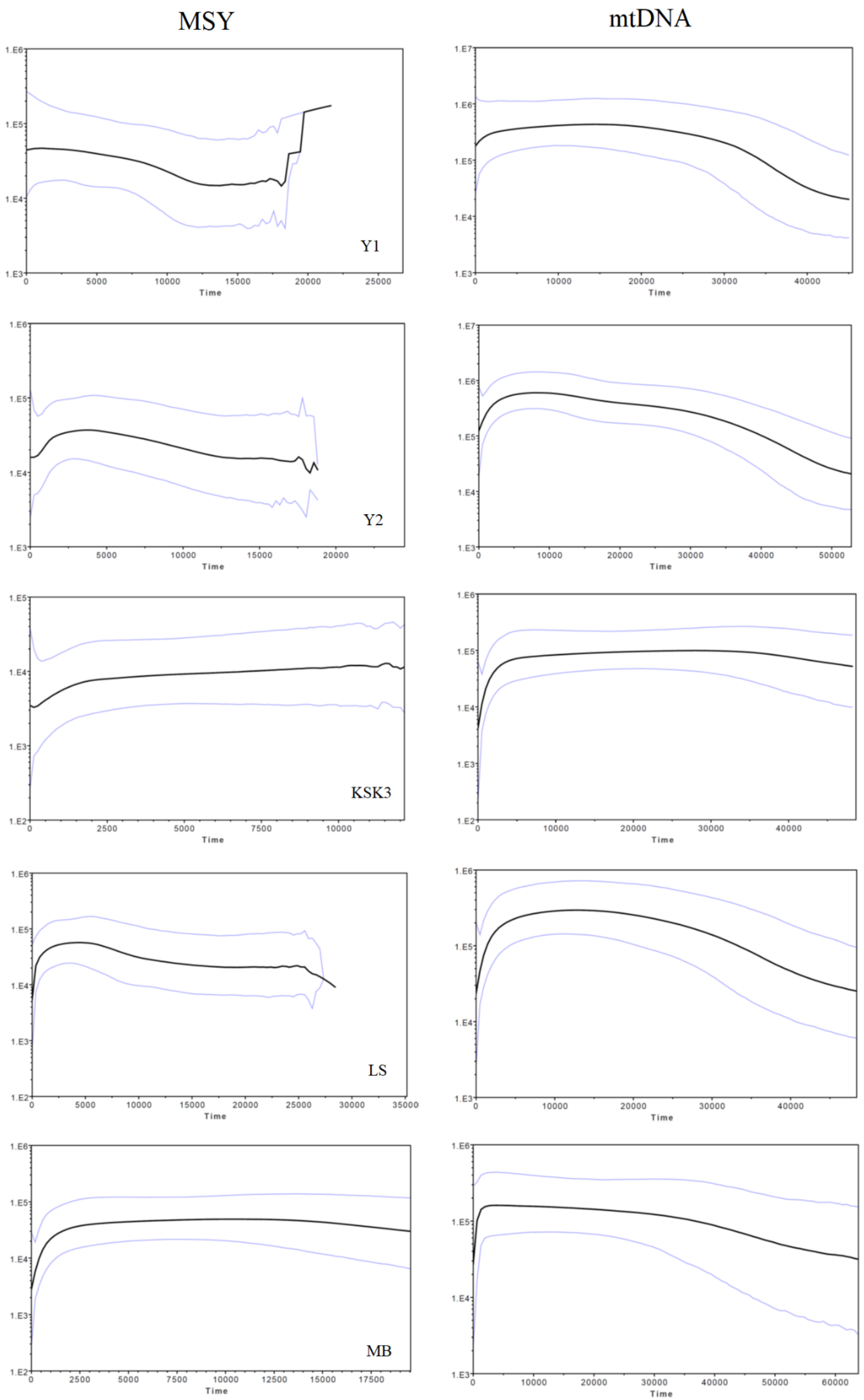
MSY
mtDNA
Y1
Y2
KSK3
LS
MB
Time

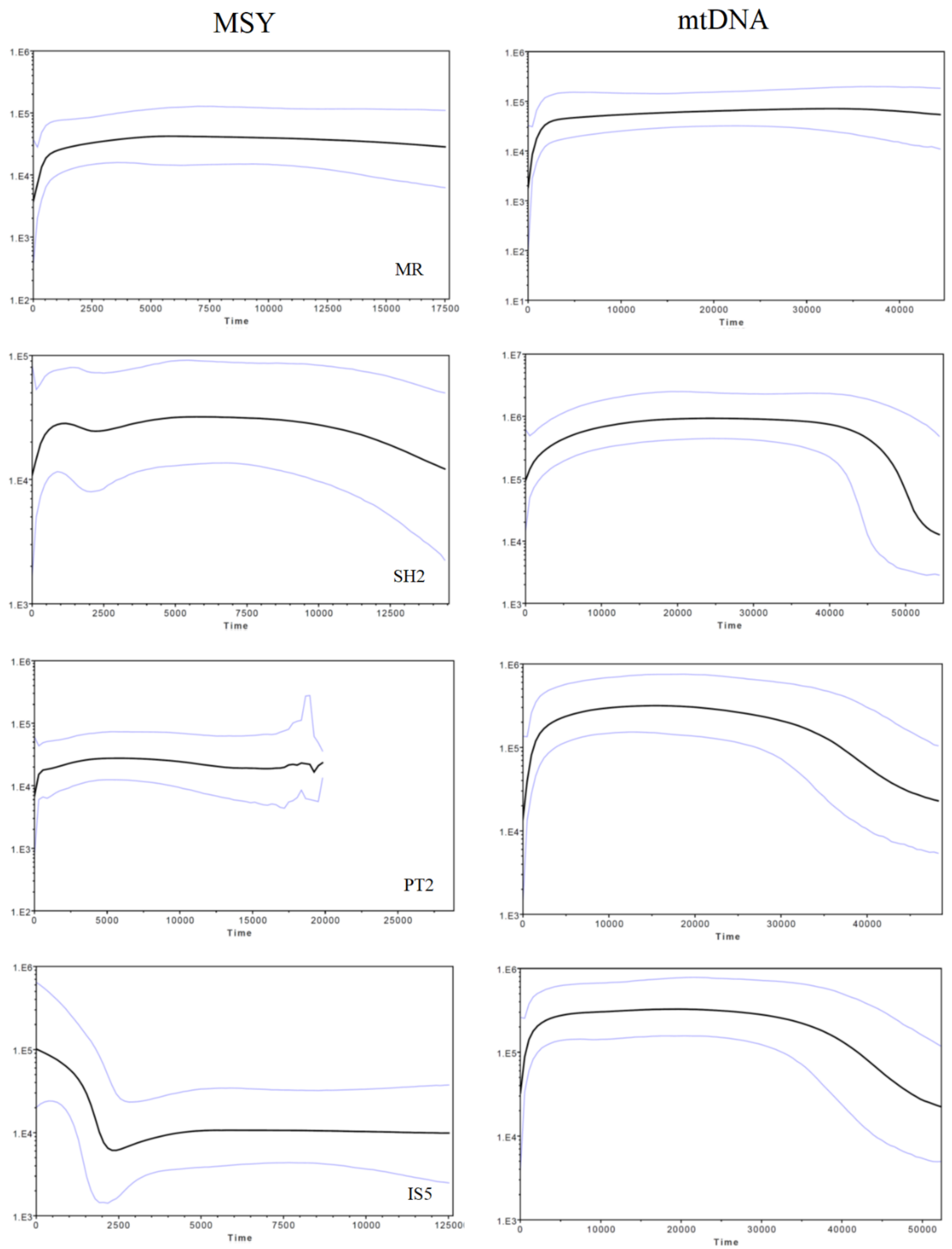


**Figure S8** The BSPs based on the MSY and mtDNA for 14 populations. Solid lines are the median estimated effective population size (y axis) through time from the present in years (x axis). The 95% highest posterior density limits are indicated by light-purple lines.

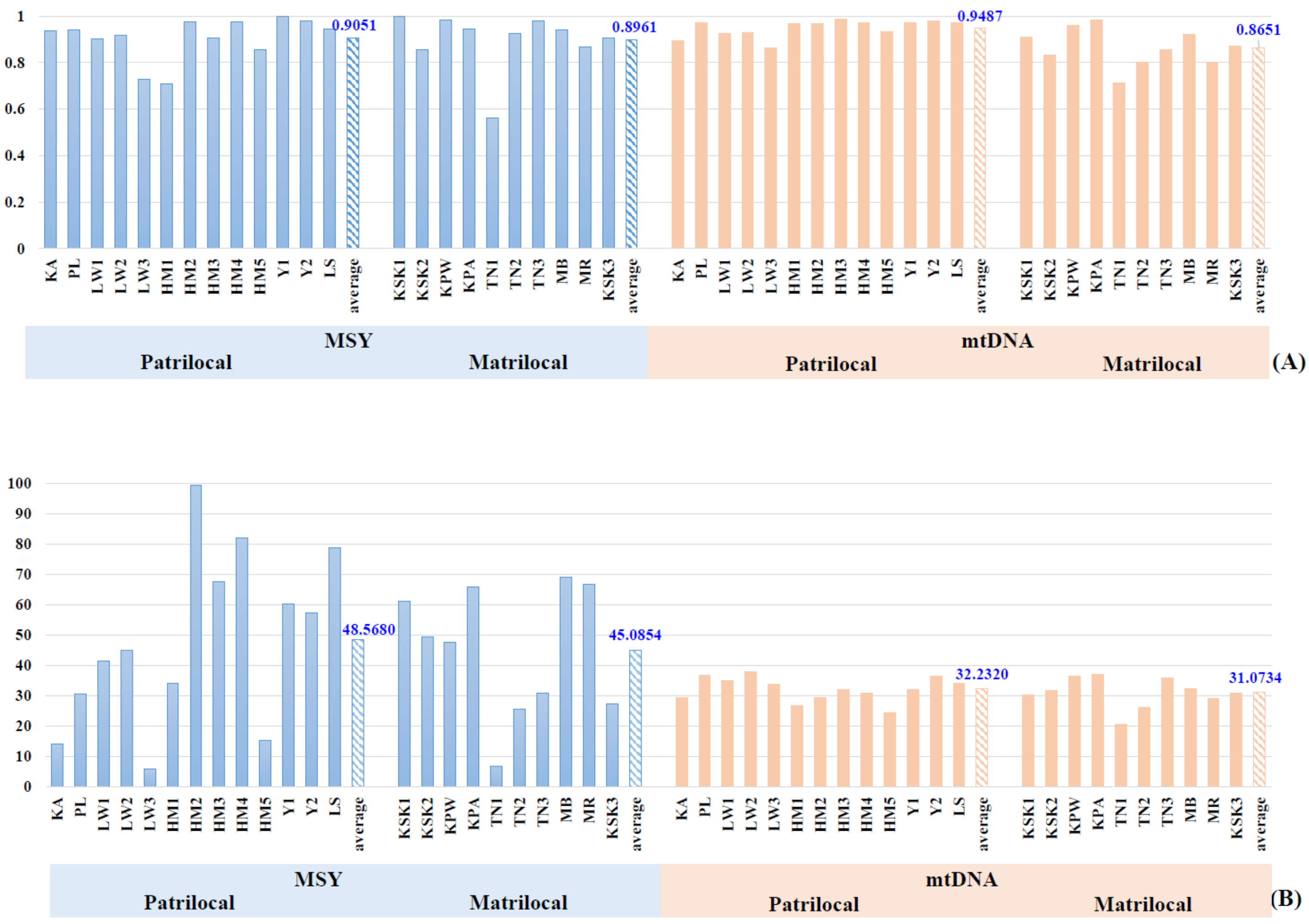


**Figure S9** Bar plots of (A) haplotype diversity and (B) mean number of pairwise differences of patrilocal (blue) and matrilocal (orange). Population abbreviations are in Table S1.

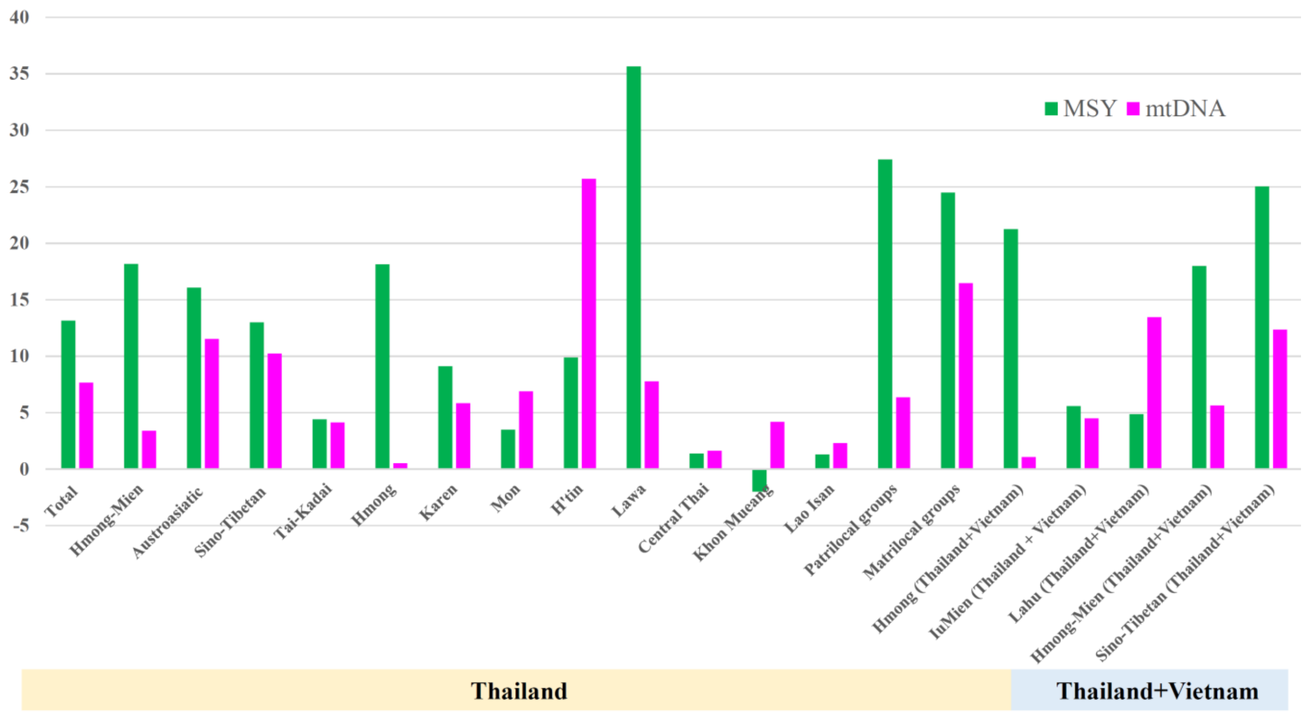


**Figure S10**. Genetic variation among populations within groups, defined by ethnicity, language and cultural practice.
